# Integrative analysis of RNA binding proteins identifies DDX55 as a novel regulator of 3’UTR isoform diversity

**DOI:** 10.1101/2025.05.06.652471

**Authors:** Matthew R. Gazzara, Timothy Cater, Michael J. Mallory, Yoseph Barash, Kristen W. Lynch

## Abstract

The 3’ untranslated regions (3’UTRs) of mRNAs play a critical role in controlling gene expression and function because they contain binding sites for microRNAs and RNA binding proteins (RBPs) that alter mRNA stability, localization, and translation. Most mRNA 3’ ends contain multiple polyadenylation sites (PAS) that can be utilized in condition-specific manners, a process known as alternative polyadenylation (APA), however the mechanisms driving the regulation of APA remain poorly characterized. By integrating a large set of over 500 RNA binding protein (RBP) depletion and binding experiments across two cell lines generated by the ENCODE consortium, we uncovered a number of RBPs in each cell type whose depletion leads to widespread alteration of 3’ UTR patterns. These include not only known regulators of APA, but also many putative novel regulators of 3’UTR isoform expression. We focused analysis on the largely unstudied DEAD box RNA helicase, DDX55, and validate its novel role in 3’UTR isoform regulation using molecular assays and targeted 3’ end sequencing experiments. Our findings identify DDX55 as a new regulator of APA, particularly at PAS that contain features of RNA secondary structure. Our data also suggest additional previously unrecognized regulators of 3’ UTR processing and differential stability.

## Background

For almost all human protein coding genes, termination of transcription and release of a mature mRNA to export to the cytoplasm occurs through a coupled reaction of cleavage of the nascent transcript followed by addition of a polyadenylate tail (polyA) [1,2]. The site at which this cleavage and polyadenylation occurs is called the polyA site or PAS and defines the 3’ termini of the message. Importantly, most human messages have more than one possible PAS site, and differential use of these PAS under distinct cellular conditions is called alternative polyadenylation or APA [1,2].

APA is a highly dynamic and widespread post-transcriptional regulatory mechanism that varies across physiological conditions and diseases [1,3,4]. Critically, APA results in mRNA isoforms with distinct 3’ untranslated regions (3’UTRs). 3’ UTRs typically contain binding sites for miRNAs and proteins that regulate mRNA stability and/or the efficiency of translation [3,5]. Therefore, the identity of the 3’ UTR of a message (i.e. how long or short the 3’UTR is and what sequences are encompassed) has profound impact on the duration and expression level of the encoded protein [3,5,6]. Consistently, APA plays crucial roles in cellular differentiation, proliferation, and stress responses [4,7–9]. Indeed, recent quantitative trait locus (QTL) studies have highlighted the genetic underpinnings of APA by linking 3’ end isoform usage to genetic variation and disease susceptibility [10,11]. These findings underscore the importance of APA in shaping transcriptomic diversity and suggest that genetic disruptions in APA contribute to pathological conditions.

Cleavage of nascent transcripts and subsequent addition of the polyA tail is mediated by a large multi-component assembly comprised of three main subcomplexes, CPSF (cleavage/polyadenylation specificity factor), CFI (cleavage factor 1) and CstF (cleavage stimulation factor) [1,2]. CFI binds to a UGUA sequence upstream of the cleavage site, while CstF binds to a UG-rich element downstream [1,2]. Both CFI and CstF promote recruitment of the CPSF complex to the central “PAS hexamer” element which typically has the sequence AAUAAA [1,2]. The CPSF complex contains the nuclease that cleaves the transcript at a CA dinucleotide 10-20 nucleotides after the PAS hexamer, and also recruits the polyA polymerase (PAP) to this location for polyadenylation after cleavage. The efficiency of CPSF activity at a particular location is dependent on the match of the PAS hexamer to consensus and the presence of adjacent CFI and CstF complexes. CFI and CstF occupancy at their cognate sites is, in turn, dependent on affinity of the binding site and the abundance of the RNA-binding subunits of CstF and CFI, namely CstF-64 and Nudt21, respectively [1,2,12].

Most well-documented examples of APA regulation have thus far been attributed to changes in the abundance of core cleavage and polyadenylation (CPA) machinery proteins, particularly CstF-64 and Nudt21 (also called CFIm25) [1,8]. However, auxiliary RNA binding proteins (RBPs), such as hnRNP H/F [13], CELF2 [14], NOVA [15], and TIA1 [16], have been shown in some systems to regulate APA. For some of these RBPs, regulation of APA has been clearly shown to be due to overlapping and competitive binding with CPA proteins, thereby blocking use of particular PAS [14]. In other cases [16] the apparent APA is more indirect, resulting from differential mRNA stability due to binding sites in the unique region of the 3’UTR.

Advances in deep learning have enabled the prediction of polyadenylation sites (PAS), improving our understanding of APA regulation [17]. However, these models remain limited in their interpretability and their ability to capture condition-specific APA patterns and sequence features beyond the well-characterized CPA motifs and factors. It is also likely that many more regulators of APA exist than have been thus far characterized, regulators who may help explain the extensive variability in 3’ UTR isoform expression across cell types and disease states. However, to our knowledge, a large-scale investigation to identify RBP regulators of APA has not been reported.

The ENCODE consortium has generated a large data set of experiments to study the roles of RBPs in gene regulation, including hundreds of depletion experiments (shRNA knockdown, CRISPRi) followed by bulk RNA-seq in HepG2 and K562 cells and RNP binding data from eCLIP experiments in these same cell types [18]. These data have been used to characterize various aspects of pre-mRNA processing including splicing, allele specific binding, and steady-state mRNA expression levels [18,19]. Here, we leverage the ENCODE data to identify new regulators of 3’UTR identity. Specifically, by integrating across over 500 RNA binding protein (RBP) depletion and binding experiments, we uncovered a number of RBPs in each cell type whose depletion leads to widespread alteration of 3’ UTR patterns. Our analysis identifies many of the known regulators of APA, supporting the relevance of our approach. In addition, we identity several putative new regulators of APA. These putative new APA regulators are RBPs assayed by ENCODE which were not previously implicated in changes to 3’UTR isoform expression. Using molecular assays and targeted 3’ end sequencing experiments, we validate one of these proteins, the DEAD box RNA helicase DDX55, as a direct regulator of APA, particularly at PAS that contain features of RNA secondary structure. Together our data expand our appreciation for the breadth of regulation of APA and 3’ UTR identity, and demonstrate at least one previously unappreciated regulatory mechanism by which 3’ UTR identity is determined.

## Results

### Analysis of hundreds of RBP knockdowns defines known and novel regulators of 3’UTR isoform expression in human cells

The ENCODE consortium has generated a large data set of results from experiments to study the roles of RBPs in gene regulation, including hundreds of depletion experiments (shRNA knockdown, CRISPRi) followed by bulk RNA-seq in HepG2 and K562 cells [18,19]. This data, together with *in vitro* motif data and *in vivo* RBP binding from eCLIP, has been used to characterize various aspects of pre-mRNA processing, including alternative splicing, allele-specific binding, and steady-state mRNA expression levels [18–20]. However, it has not, to our knowledge, been used to characterize the landscape of alternative polyadenylation (APA) and 3’UTR isoform diversity. Therefore, we first sought to identify the functional APA targets of RBPs by comparing control to depletion conditions to identify target genes of APA regulators.

Accurate quantification of APA from bulk RNA-seq data is challenging and numerous tools have been developed for this task ([17] and references therein). We and others have benchmarked popular tools using different synthetic and orthogonal datasets as ground truths [17]. While each tool has unique strengths and weaknesses, we chose to focus our analysis using the highly utilized DaPars algorithm [21] to call tandem APA changes upon RBP knockdown. Previous work in our lab established thresholds that yield high validation rates using DaPars [14,22]. Additionally, the fact that DaPars can identify polyadenylation sites (PAS) *de novo* from RNA-seq read coverage and is not limited to just annotated PAS was appealing for our analysis where RBP knockdown could cause creation of previously unused PAS absent in the annotation.

We uniformly processed the available RBP depletion experiments from ENCODE (235 RBPs in HepG2, 244 RBPs in K562; Supplementary Table 1,2) with DaPars and called significant changes in APA as genes containing a terminal exon with an absolute shift in PAS usage greater than 20% (change, or delta, in Percentage of Distal polyA site Usage Index, denoted|dDPUI|>20%)) with a false discovery rate of less than 5% (see Methods). We observe a large number of gene terminal exons with significant tandem APA shifts upon RBP knockdown in both HepG2 (Fig 1A, purple), and K562 cells (Fig S1, purple). Several of the RBPs with the highest number of targets have previously been implicated in regulation of 3’end processing or other aspects of 3’UTR isoform regulation (Fig 1A, S1; asterisks), but many others have not previously been identified as 3’end regulators. Interestingly, the RBPs with the greatest number of APA changes in HepG2 and K562 cells were DBHS family proteins NONO and SFPQ (also known as PSF), respectively, which alter over one thousand target genes each. Both proteins have been shown to regulate APA [3,23–25] and a number of other molecular processes [23,26].

**Fig 1:**
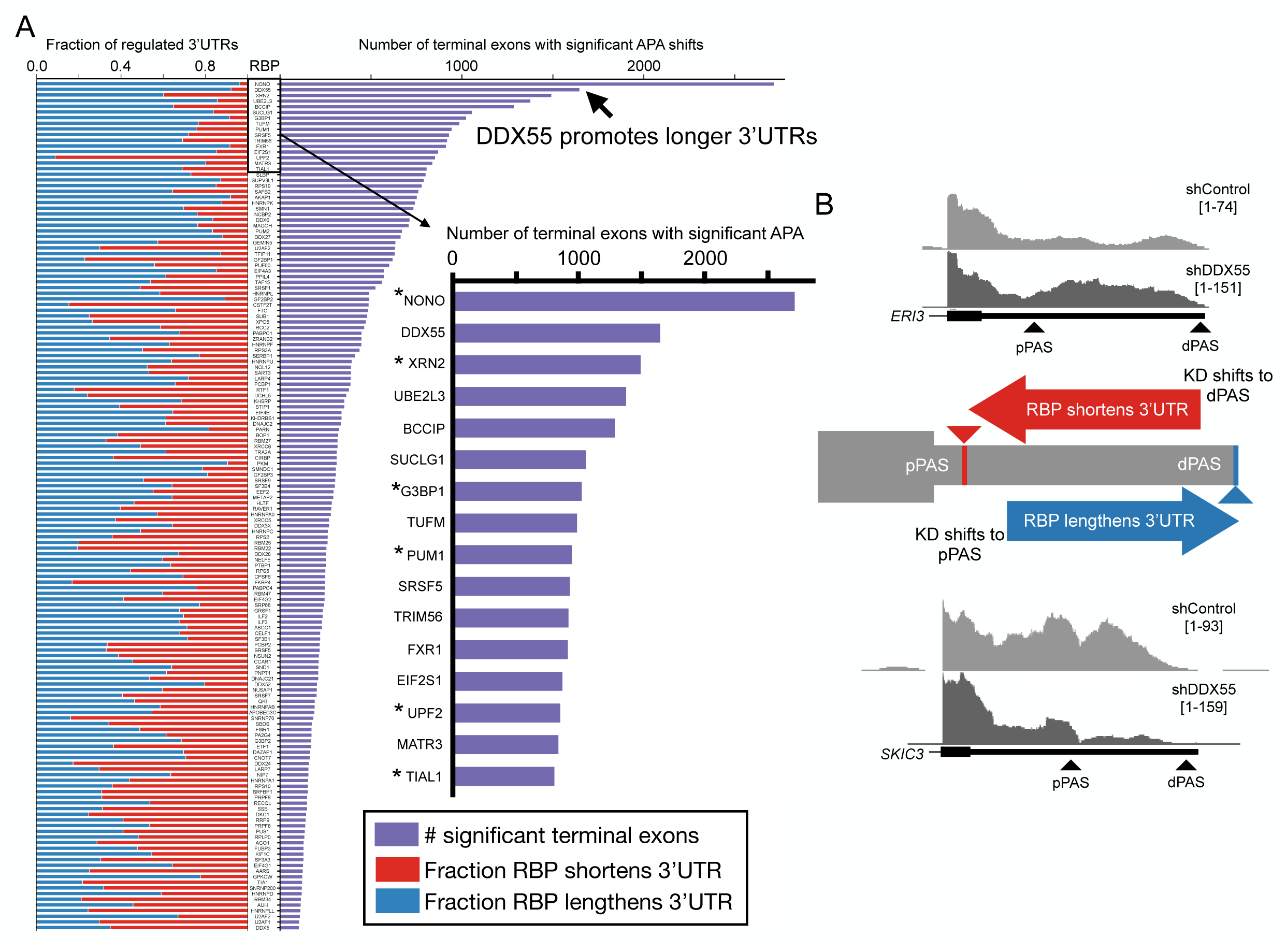
The landscape of 3’UTR tandem APA regulation in HepG2 cells. (**A**) DaPars analysis of top 150 HepG2 RBP knockdowns from ENCODE (all 235 experiments listed in Supplementary Table 1) sorted by the number of significant terminal exons (right, purple, |dDPUI|≥20% with adjusted *p*<0.05). Right inset shows the top RBPs by number of terminal exons regulated with RBPs known to influence 3’UTR isoforms indicated by asterisks. Stacked bar chart on the left indicates the fraction of significant terminal exons that had a pattern of RBP promoting shorter 3’UTRs (red, left bars) or RBP promoting longer 3’UTR (blue, right bars). (**B**) Center panel shows a cartoon illustration of two classes of terminal exon (TE) tandem APA shifts that emerge from ENCODE RBP data: TEs where the RBP normally promotes pPAS usage/short 3’UTR expression (knockdown shifts coverage towards dPAS, top, red) or TEs where the RBP promotes dPAS usage (knockdown shifts coverage towards pPAS, bottom, blue). Top panel shows UCSC genome browser RNA-Seq coverage example of a terminal exon in *ERI3* that is normally shortened by DDX55 (KD, bottom, shifts to dPAS). Bottom panel shows example of a terminal exon in *SKIC3* (also known as TTC37) that is normally lengthened by DDX55 (knockdown, bottom, shifts to pPAS).

Tandem APA shifts in terminal exons sometimes show biases toward 3’UTR lengthening or shortening in specific conditions or disease states. For example, multiple studies suggest proliferative cell states tend to express shorter 3’UTRs and more differentiated cell states express longer 3’UTRs conditions [27–29]. This has notable and particularly important implications for certain cancers where increased expression of core CPA factors (e.g. CstF-64/CstF-64t) drives shifts towards weaker, proximal polyadenylation sites (pPAS) [29–31]. Given this, we monitored the ENCODE data to identify classes of events regulated by each RBP. Cases where the RBP typically promotes shorter 3’UTR formation are noted in red (Fig 1A-B, e.g. top RNA-seq coverage tracks in 1B) while instanced in which the presence of the RBP promotes longer 3’UTR expression are in blue (Fig 1A-B, e.g. bottom RNA-seq coverage tracks in 1B). Across the hundreds of RBP knockdowns we find a range of RBPs that have a bias towards promoting cleavage and polyadenylation at the distal site (dPAS) to generate long 3’UTR expression (e.g. DDX55, Fig 1A, S1) or promoting pPAS to shorten 3’UTR expression (e.g. UPF2, Fig 1A, S1). Additionally, the isoform shifts we observe upon knockdown of the core CPA component CstF-64/64t (encoded by *CSTF2/2T* respectively) are consistent with previous results where co-depletion of these factors in HeLa cells resulted in general 3’UTR lengthening, suggesting these proteins promote usage of weaker pPAS that rely on their enhancing effects [32]. Taken together, this suggests our analysis of ENCODE RNA-seq data can recapitulate known trends in APA regulation by RBPs.

Given that NONO and the RNA Pol II termination factor and exonuclease XRN2 are established regulators of APA [3,25], we were particularly intrigued by DDX55 since it regulates a similar number of 3’UTRs as these known regulators in HepG2 cells (over 1,500 terminal exons). Of note, genes regulated by DDX55 exhibit a striking pattern of preferential DDX55-dependent lengthening of 3’UTRs (Fig 1A, >95%). Manual inspection of read coverage tracks of top putative targets of DDX55 showed clear shifts in PAS usage from control to knockdown cells (Fig 1B, top and bottom). There is a similar trend in K562 cells for DDX55 depletion to cause 3’UTR shortening (Fig S1), although we observe fewer DDX55-dependent APA shifts (around 250 terminal exons) in K562 cells than HepG2, perhaps owing to the less efficient DDX55 knockdown in the K562 cells (45-60% depletion in K562, 66-70% depletion in HepG2 by Western blot: www.encodeproject.org).

DDX55 is a member of the DEAD-box helicase family of proteins (Fig S2A) and is an essential gene (Fig S2B) which is broadly expressed across different tissues (Fig S2C). DDX55 is localized in the nucleus as well as the cytoplasm in several cell types [33,34]. Some cell types also show DDX55 localization to nucleoli [34,35], consistent with a demonstrated role of DDX55’s yeast homolog Spb4 in rRNA processing [36]. In HepG2 cells we observe DDX55 primarily in the nucleus (Fig S2D), consistent with DDX55 impacting APA. Others have confirmed nuclear localization of DDX55 in HepG2 cells and further find DDX55 to be excluded from the nucleolus in these cells [33].

### eCLIP analysis identifies DDX55 as a common mRNA 3’UTR binding protein that associates near sites of cleavage and polyadenylation

Next, we turned to available eCLIP data (103 RBPs in HepG2 and 120 RBPs in K562, Supplementary Table 3) to identify which RBPs bind within 3’UTRs and thus may directly influence 3’UTR isoform expression via APA and/or other mechanisms such as mRNA stability. We first defined expressed transcripts as those with TPM≥1 in both cell types. We then identified RBPs with any evidence of binding (i.e. presence of any eCLIP peak) in a 3’UTR and sorted for the top 3’UTR binders in HepG2 cells (Fig 2A) and K562 cells (Fig S3A). DDX55 ranked near the top of both lists (4th in HepG2, 2nd in K562) and bound ∼80% of expressed 3’UTRs. Other pervasive 3’UTR binders included RPBs with known functions in 3’end processing and/or mRNA stability such as LARP4 [37], IGF2BP1/3 [38], TIA1/TIAL1 [16], or UPF1 [39]. For each RBPs we also determined what percentage of high-confidence peaks fall within 3’ UTRs versus other genomic regions across genes expressed in HepG2 (Fig 2B). Once again, DDX55 ranked near the top of the list (4th), with over 80% of its binding events occurring within expressed 3’UTRs. Other annotated features bound by DDX55 include CDS exons and non-coding exons (Fig 2C).

**Fig 2:**
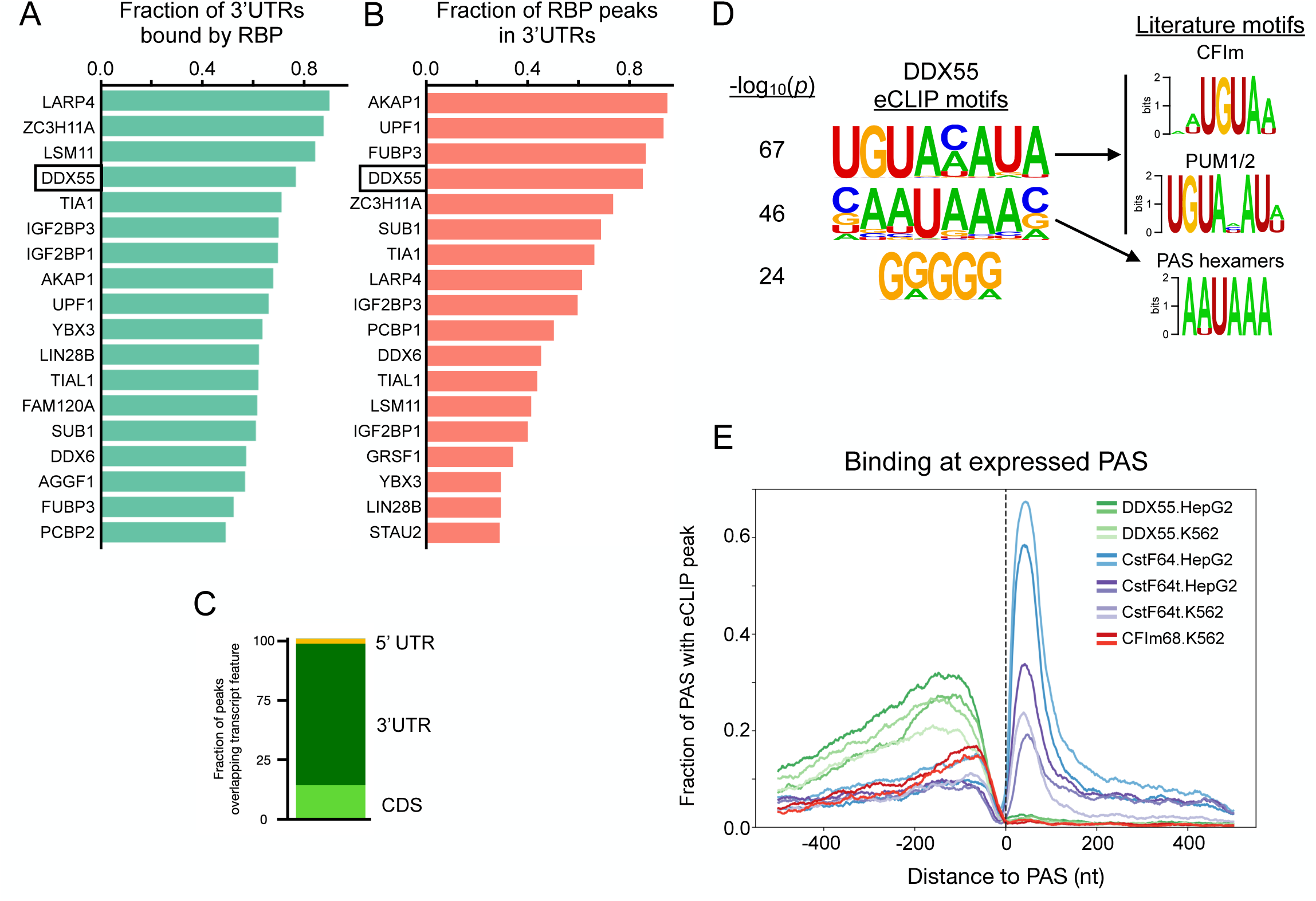
The landscape of 3’UTR binding RBPs identifies DDX55 as a pervasive 3’end binding protein. (**A**) Fraction of expressed mRNA transcripts (TPM≥1) from HepG2 cells that had evidence of RBP binding in HepG2 cells (presence of any eCLIP peak) within the 3’UTR. (**B**) Fraction of high confidence HepG2 eCLIP peaks based on IDR (Irreproducible Discovery Rate) for each RBP which overlapped with 3’UTRs of expressed mRNA transcripts (TPM≥1 in HepG2). (**C**) Fraction of DDX55 peaks from HepG2 cells that overlapped each transcript feature. (**D**) Top enriched motifs discovered from sequences covered by HepG2 DDX55 IDR peaks using HOMER (left) and matches to literature motifs for the CFIm complex (motif for CFIm25, encoded by *NUDT21*, shown) [12], the Pumilio Response Element that binds PUM1/2 to influence mRNA stability [67], or the top two know PAS hexamer signals (AAUAAA or AUUAAA) (right). (**E**) RNA map of the fraction of expressed PAS from human liver with evidence of DDX55 binding (greens) or core CPA machinery binding (blues, purples, reds) in indicated cell type with two replicate eCLIP experiments per condition.

Core sequence motifs are known to regulate 3’UTR isoform diversity through the binding of the CPA machinery or the recruitment of proteins to alter mRNA stability. Given DDX55’s pervasive and specific binding profile to 3’UTRs, we calculated enriched motifs under high confidence peaks. Surprisingly, we found the most significant motifs to match the known binding sites for the core CPA complex CFIm (UGUA) and the Pumilio Response Element that binds PUM1/2 (UGUAHAUA) (Fig 2D). The second most significant motif was the core PAS hexamers (AAUAAA / AUUAAA) (Fig 2D). This suggests DDX55 binds near functional elements in 3’UTRs that influence 3’end formation through cleavage and polyadenylation.

Given the match of DDX55’s top eCLIP motifs to some of the core CPA signals, we first examined the per-nucleotide binding profile of all RBPs in ENCODE to see which positional biases, if any, exist around the PAS that are used in human tissues, regardless of regulation. We therefore mapped CLIP-peak density for each RBP around PAS sites identified from targeted 3’ end sequencing from human liver for HepG2 eCLIP experiments or lymphocytes for K562 experiments (Fig S3B,C). CstF-64 and CstF-64t are two components of the heterotrimeric CstF complex which binds the Downstream Sequence Element (DSE) that is essential for the cleavage reaction. CLIP data exists for both of these proteins in both cell lines, so we used these proteins to set a baseline for our analysis of other RBP binding to the region. However, our initial analysis failed to give the expected positional enrichment for these known RBP binding regions (Fig S4). Upon further investigation, we realized that the CLIPper algorithm, which ENCODE uses to call eCLIP peaks, only searches for binding within the boundaries of annotated transcript 5’ and 3’ ends. Thus, the primary site of CstF binding (i.e. downstream of the 3’ end cleavage site) is often missed. To correct for this, we extended gene boundaries by 500 nt downstream and re-called peaks with CLIPper for all eCLIP experiments in ENCODE. Use of this extended annotation resulted in CstF-64/CstF-64t peaks being called over their known binding motif downstream of the annotated gene ends, which were missed in the original analysis (Fig S4). Correcting this peak calling also has important implications for other regulatory proteins that bind downstream of the 3’ cleavage site. We therefore provide our downstream-aware peak calling as a resource (https://doi.org/10.5281/zenodo.15133362) to better define RBP binding in this region and to remove potential false-positive calls upstream of the cleavage site.

We applied the aforementioned downstream-aware peak calling procedure to all RBPs in both cell types. We found FMR1 in K562 cells also binds in a tightly-localized region downstream of 3’ cleavage sites, as observed for CstF-64/64t (Fig S3C). All other proteins surveyed primarily associate with 3’ UTRs upstream of the 3’ cleavage site. We noticed that proteins known to regulate mRNA stability tend to interact with RNA in a diffuse pattern along the entire length of the 3’ UTR (e.g. IGF2BP1/3, UPF1, YBX3) [39–41], while proteins that have been shown to directly regulate the CPA machinery (e.g. TIA1 and LAPR4 [10,16]) exhibit enriched binding closer to the site of cleavage (Fig S3B-C). Of note, the association of DDX55 with 3’ UTRs also displays a bias towards the 3’end PAS (Fig S3B-C). Furthermore, plotting an RNA-map of binding for DDX55 and all core CPA proteins with eCLIP from ENCODE showed DDX55 bound proximal to as many PAS sites as core CPA factors like CFIm68 or CstF-64t (Fig 2E). Interestingly, pairwise overlap of all eCLIP experiments by Jaccard Index showed DDX55 and CFIm68 clustered separately, but both overlapped with known APA and mRNA stability regulators (Supplemental Fig S5).

### Integrating binding and RNA-seq functional targets identifies DDX55 as a direct regulator of 3’UTR isoform expression

Having confirmed that DDX55 exhibits a binding profile consistent with other 3’ end processing factors, we next wanted to ask if and how DDX55 associates around PAS sites that exhibit altered use (i.e. APA) in a DDX55-dependent manner. We therefore integrated the functional targets from DaPars with eCLIP for DDX55 targets in HepG2 cells. Strikingly, when comparing target terminal exons where DDX55 promotes 3’UTR lengthening (1518 terminal exons), we observed clear enrichment of DDX55 binding specifically in the alternative 3’UTR region, between the DDX55-inhibited proximal PAS and the DDX55-promoted distal PAS, compared to a control set of non-regulated targets (Fig 3A,B, S6A). Indeed, the strongest enrichment of DDX55 binding is just upstream of the dPAS that is favored in the presence of DDX55 (Fig 3A,B, S6A). This strong pattern of binding was not observed in cases in which DDX55 promotes use of the pPAS, although some DDX55 association with the 3’ UTR was observed in these cases (Fig 3C,D, S6B).

**Fig 3:**
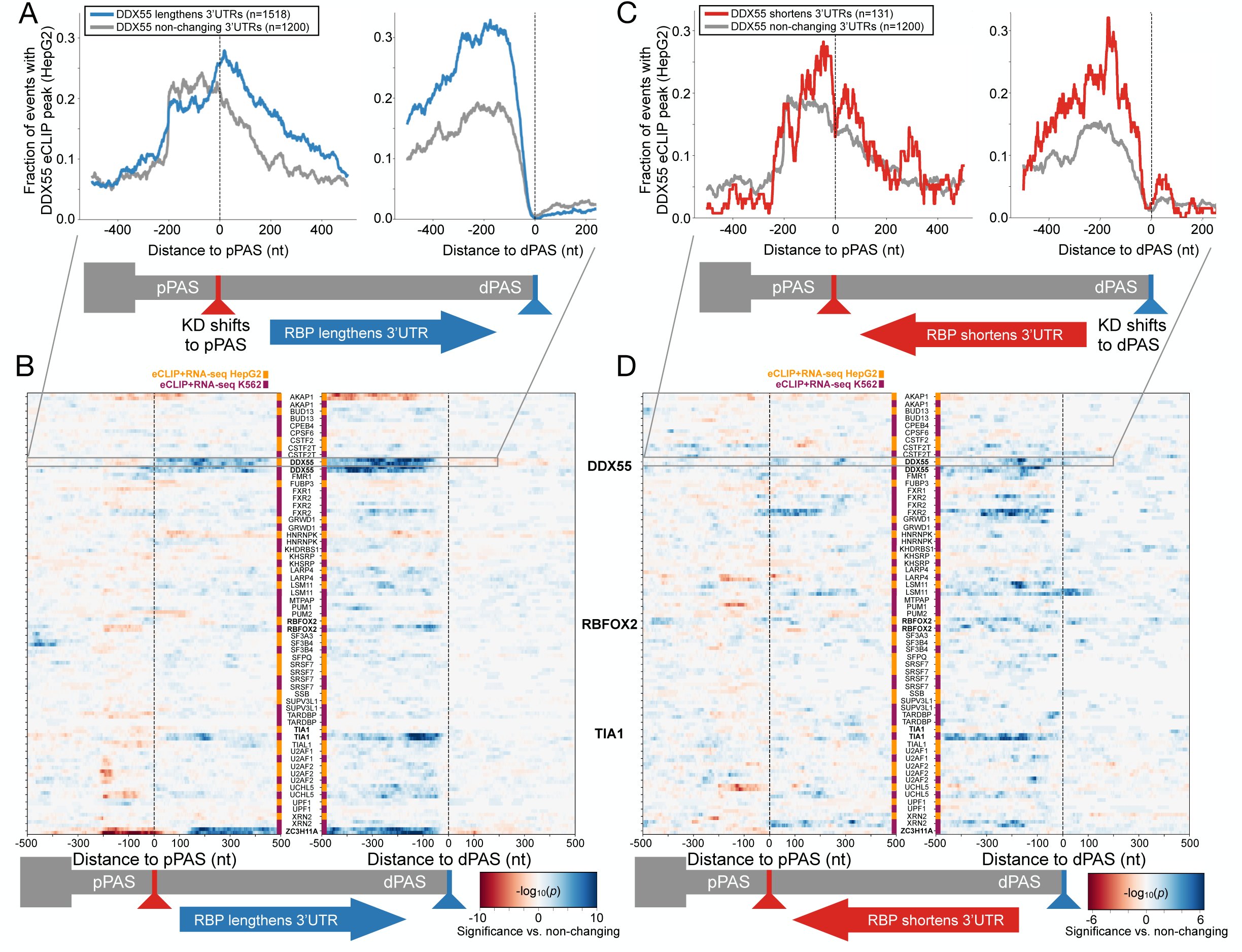
Integrating functional and binding targets of tandem APA identifies direct regulators of 3’UTR isoform expression. (**A**) RNA map of the fraction of events with HepG2 DDX55 eCLIP peaks centered around proximal polyadenylation sites (pPAS, left) or distal PAS (dPAS, right) for events where DDX55 promotes 3’UTR long isoform expression (blue line, knockdown (KD) shifts expression towards pPAS, dDPUI≥20% with FDR<0.05) or events were DDX55 depletion had no effect on PAS choice (gray line, |dPDUI|≤5% with FDR>0.05). (**B**) Heat map showing the significance of enrichment (two-tailed Fisher’s exact test -log10(*p*-value)) of RBP binding at each position proximal to regulated pPAS (left) and dPAS (right) for each RBP in ENCODE HepG2 (orange) and K562 cells (purple). Shown are all the RBPs which had both matching eCLIP and RNA-seq depletion data comparing terminal exon sets that show RBP regulated 3’UTR lengthening events (RBP KD shifts expression towards pPAS, dDPUI≥20% with FDR<0.05), versus those not regulated by the RBP. (**C**) As in A, but for events where DDX55 promotes 3’UTR short isoform expression (red line, knockdown, denoted KD, shifts expression towards pPAS, dDPUI≤-20% with FDR<0.05) compared to non-changing events (grey). (**D**) As in B, but for RBP 3’UTR shortening events (RBP KD shifts expression towards dPAS, dDPUI≤-20% with FDR<0.05) versus those not regulated by the RBP.

We extended this analysis to all RBPs in ENCODE which had an RNA-seq depletion experiment and eCLIP binding data within the same cell line (88 RBPs in HepG2, 119 RBPs in K562), and calculated the significance of binding enrichment or depletion for events where the RBP normally lengthens (Fig 3B) or shortens the 3’UTR (Fig 3D), versus non-regulated genes. Compared to all RBPs in ENCODE, DDX55’s binding profile stands out as one of the most enriched over the targets it regulates, particularly for the large class of 3’UTRs where it promotes long 3’UTR formation (Fig 3B). This enrichment was significant in both cell types (Fig 3B, S6-S8), despite K562 cells having fewer DDX55 responsive targets (Fig S1). Indeed, the pattern of DDX55 binding around PAS sites regulated in a DDX55-dependent manner mirrors that of other RBPs, such as TIA1 and RBFOX2 (Fig 3B, S7), which have been shown to regulate 3’ UTR identity [16,42].

Interestingly, several well-documented APA regulators did not have strong positional enrichment in this analysis, suggesting a lack of specific location from which they function or indirect activity. Some we could not analyze since matching eCLIP and knock-down experiments in the same cell type do not exist (e.g. the DBHS proteins NONO in HepG2 and SFPQ in K562). Others, like the exonuclease XRN2, may be non-specific binders and/or be difficult to capture since they are associated with actively degrading the RNA. We also note interesting enrichment profiles of other select RBPs that may be playing a direct role in short 3’UTR isoform choice (e.g. IGF2BP1, PCBP2; Fig S8). In sum, the binding profile of DDX55 along 3’UTRs is consistent with a previously unrecognized activity of DDX55 in directly regulating 3’ UTR identity, likely through direct influence on the CPA machinery.

As discussed above, 3’ UTR identity is initially determined by the site of cleavage and polyadenylation in the nucleus (APA, Fig 4A, left). However, since the 3’UTR can influence the stability of the mRNA in the cytoplasm, differential stability conferred by long or short 3’ UTRs is also a mechanism by which preferential 3’ UTR abundance may be observed in the cytoplasm (Fig 4A, right). Therefore, to further investigate the mechanism by which DDX55 impacts 3’ UTR identity we carried out targeted 3’ end sequencing of RNAs isolated specifically from the nucleus or cytoplasm. We confirmed both the efficiency of DDX55 depletion in both compartments, as well as the integrity of the fractionation, by western blot (Fig 4B), isolated RNA from both fractions and carried out targeted 3’end sequencing (see Methods) to specifically map the 3’ termini of mRNAs (i.e. sites of polyadenylation) and identify changes in the distribution of 3’ termini in DDX55-depleted cells versus normal controls. Importantly, the vast majority of the DDX55-dependent changes we observe in the cytoplasm are also changing in the nucleus, consistent with a mechanism of APA during transcription (Fig 4C). This includes over 200 genes which exhibit significant changes in 3’UTR length in both the cytoplasm and nucleus, including 3’UTRs that are both lengthened and shortened upon depletion of DDX55 (Fig 4C). By contrast, we identified only 12 genes for which there was a greater change in 3’ UTR abundance in the cytoplasmic mRNA as compared to nuclear mRNA (Fig 4D). Given the limited number of 3’UTR changes that are only observed in cytoplasmic fractions and the nuclear-biased distribution of DDX55 in HepG2 cells, we conclude that DDX55 shapes 3’ UTR identity primarily through direct regulation of APA. However, we acknowledge that we can not fully rule out that in some conditions or cell-types DDX55 may act also in controlling mRNA stability through binding to specific 3’ UTRs.

**Fig 4:**
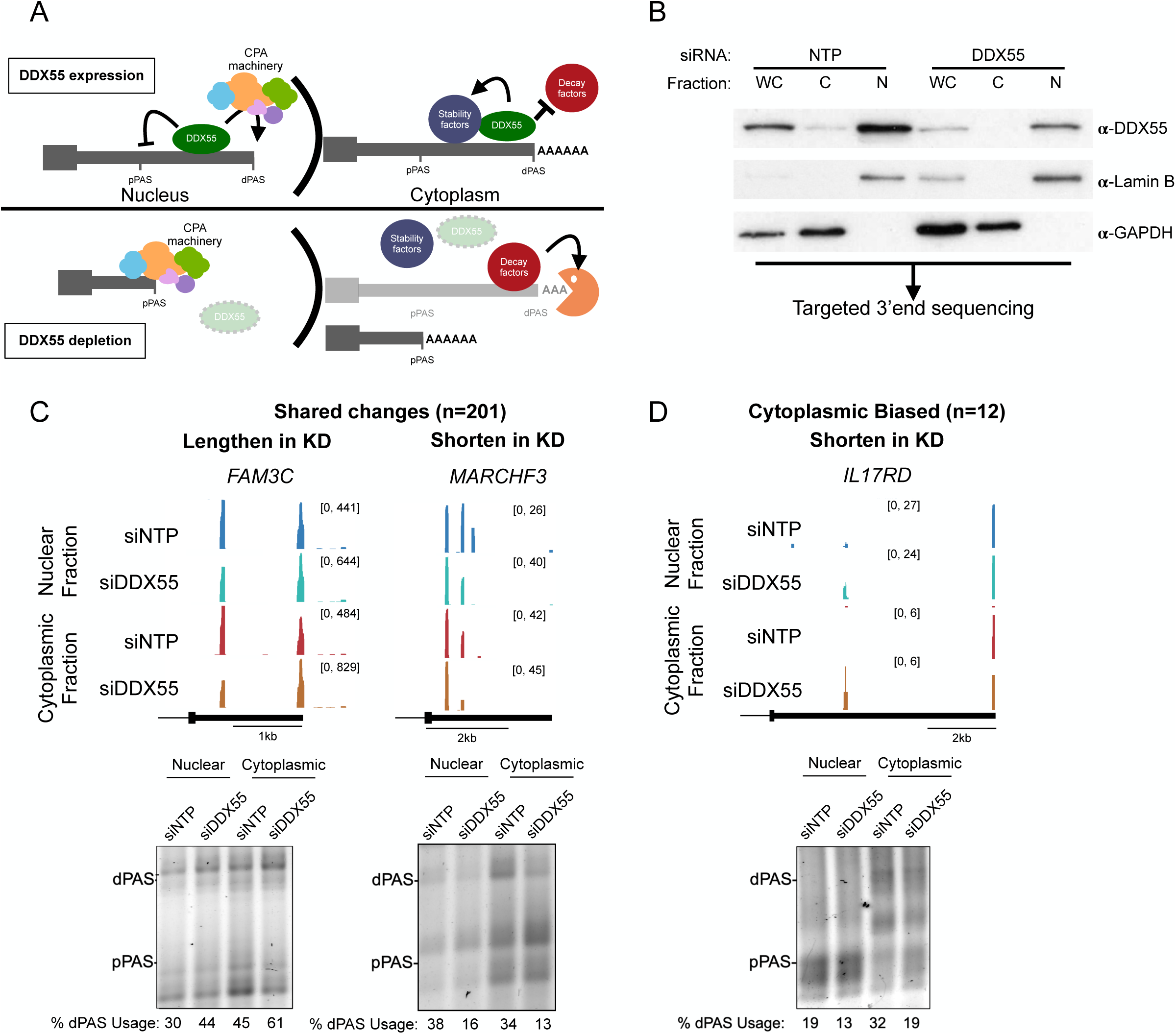
DDX55 regulates 3’UTR isoform choice in both the nucleus and the cytoplasm. (**A**) Hypothesized models for 3’UTR isoform shortening observed upon DDX55 depletion in the nucleus (left) and/or cytoplasm (right). (**B**) Western blot on protein lysates from whole cell (WC), cytoplasmic (C), and nuclear (N) fractions from HepG2 cells transfected with siRNA against a control non-targeting pool (NTP, left 3 lanes) or against DDX55 (right 3 lanes). GAPDH and Lamin B served as controls for cytoplasmic and nuclear fractionation, respectively. Total RNA from each condition was subjected to QuantSeq Rev V2 targeted 3’end sequencing. (**C**) UCSC genome browser example views of QuantSeq Rev V2 3’end clusters for DDX55 regulated termini that are changed in both the nucleus and cytoplasm (top), with 3’ RACE validations below (bottom). (**D**) same as (C) but for the few genes that exhibit changes in 3’ termini that are biased towards the cytoplasmic fraction.

### DDX55-regulated PAS are marked by features suggesting RNA secondary structure

Finally, to gain an understanding of how a helicase such as DDX55 might influence PAS choice and regulate APA, we investigated the sequence features that correlate with DDX55 binding and regulation. Most helicases bind to, and unwind, double-stranded or structured RNAs rather than associating with specific nucleotide sequences [43]. However, as shown above (Fig 2D), DDX55-associated sequences encompass sequence motifs that direct the core CPA machinery. We reasoned, therefore, that DDX55 might bind to PAS sites with particular structures. For typical PAS, the UGUA enhancer is located 40-60 nucleotides upstream of the AAUAAA hexamer [2,12]. Indeed, we observe this spacing in sequences bound by CFIm68, a component of the CFI complex of the CPA (Fig 5A). By contrast, the same analysis of sequences bound by DDX55 reveals an extended spacing of ∼120 nucleotides between the UGUA and AAUAAA, which flank either side of the center point of DDX55 association (Fig 5B). The tendency for more distance between the UGUA and AAUAAA motifs in DDX55-bound regions compared to all CFIm68-bound sites is recapitulated in both cell types tested (Fig 5B, S9A,B top) and also when the analysis is restricted to bona fide PAS (Fig 5C, S9C). However, interestingly, when we further subset the data, the extended spacing of UGUA and AAUAAA is only observed for PAS that are preferentially used upon depletion of DDX55 (e.g. the proximal PAS) and not in the DDX55 permissive dPAS (Fig S9D).

**Fig 5:**
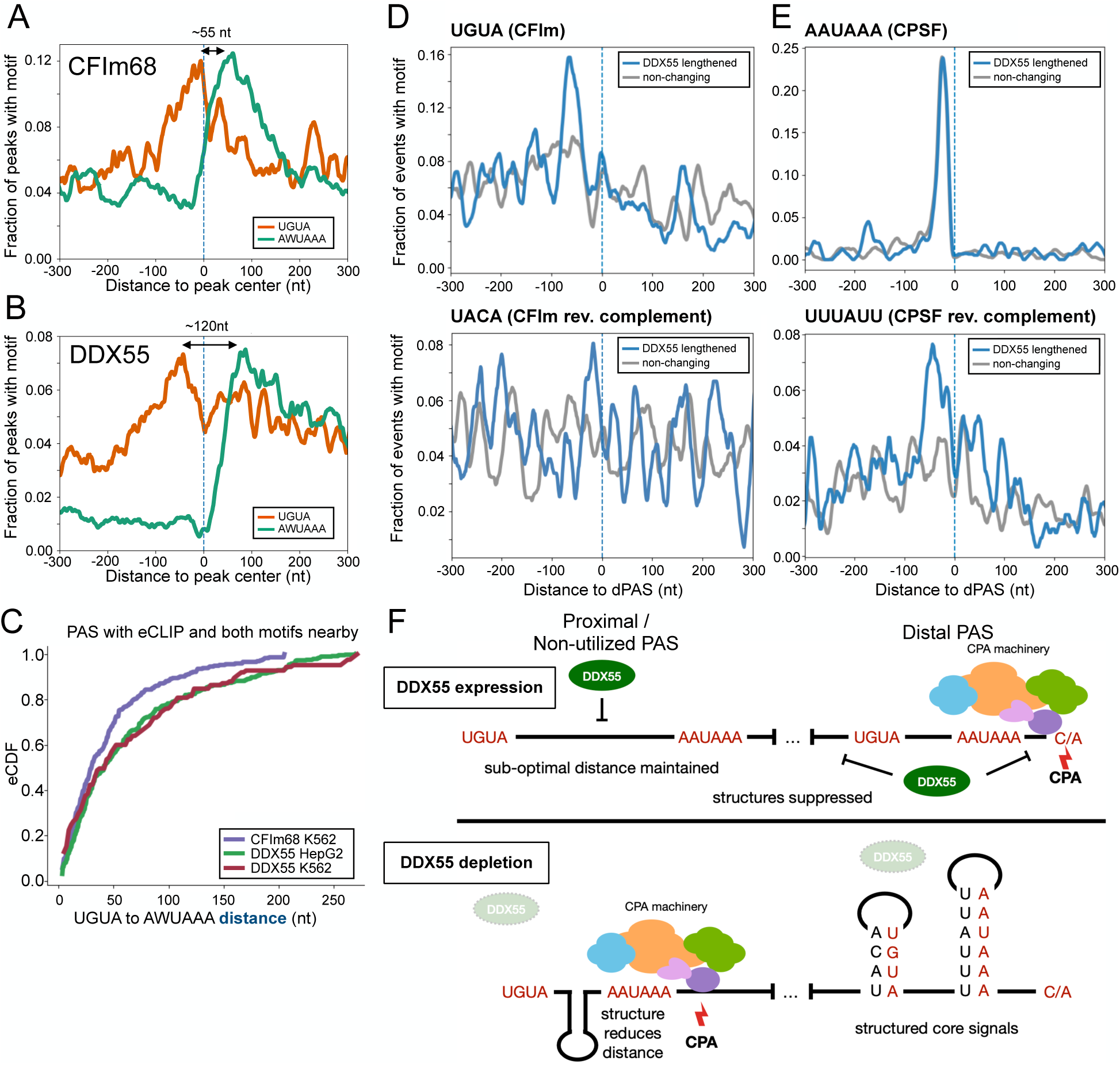
Sequence features of DDX55 and CFIm binding suggests mechanisms of regulation. (**A**) Motif maps showing per nucleotide frequency of the top two core PAS hexamers (AAUAAA or AUUAAA, green) and CFIm upstream enhancing element (UGUA, orange) centered on the high confidence eCLIP peak centers near PAS (see Methods) for CFIm component CFIm68 (CPSF6) in K562 cells. (**B**) Same as (A) but for eCLIP peak centers for DDX55 in HepG2 cells. DDX55 in K562 cells is shown in Fig S9. (**C**) CDF showing the distance between CFIm and PAS hexamer motifs within 150 nt of PolyA_DB4 sites that have proximal binding (within 100 nt) of CFIm68 in K562 cells (purple), DDX55 in HepG2 (green), or DDX55 in K562 cells (red). (**D**) Motif maps showing per nucleotide frequencies of the CFIm upstream enhancing element (UGUA, top) or its reverse complement (UACA, bottom) around distal PAS (dPAS) regulated by DDX55 (blue) or non-changing (gray). (**E**) Same as D, but for the core PAS hexamer (AAUAAA, top) or its reverse complement (UUUAUU, bottom) around distal PAS (dPAS) regulated by DDX55 (blue) or non-changing (gray). (**F**) CDF showing distance between AAUAAA and its reverse complement UUUAUU around indicated DDX55 regulated (blue) or non-regulated (gray) dPAS. (**G**) Model for DDX55 regulation of PAS choice. When expressed, (top) DDX55 suppresses PAS usage of upstream PAS with non-ideal motif distances and promotes usage of a subset of distal sites which have nearby reverse complement sequences and are structured. When depleted (bottom), sequences normally bound by DDX55, including potential PAS with longer linker regions between UGUA and PAS hexamers are folded and come closer together. Other sites with shorter linker regions at strong distal sites with reverse complements nearby are folded in the absence of DDX55.

To further investigate the sequence context of DDX55-sensitive PAS, we assessed the sequences between the UGUA and AAUAAA of the DDX55-repressed pPAS. In HepG2 cells we find that the intervening sequence between the UGUA and AAUAAA at DDX55-bound PAS sites is highly GC-rich (Fig S9A, bottom). In K562 cells the entire region around the DDX55-bound PAS is highly GC-rich (Fig S9B, bottom). By contrast, the primary feature of PAS that are preferentially utilized in the presence of DDX55 (e.g. dPAS) was the presence and proximity of sequences that are complementary to the core UGUA and AAUAAA motifs (Fig 5D-E). Such complementary sequences are not observed near DDX55-repressed pPAS (Fig S9E). Taken together, this sequence analysis suggests a model in which DDX55 may repress pPAS by unwinding structures between the UGUA and AAUAAA to maintain suboptimal spacing (Fig 5F). Additionally, or alternatively, DDX55 may promote the use of dPAS by preventing the base-pairing of critical signals (i.e. UGUA and AAUAAA) with neighboring regions of the RNA, which would be predicted to sequester access of these cis-regulatory signals by the CPA machinery. Either model explains how a protein with limited sequence-specificity such as DDX55 may regulate specific patterns of APA.

## Discussion

### An atlas of RBPs that bind and influence 3’ UTR expression to facilitate future research on mechanisms of 3’ UTR regulation

Despite the well-documented extent of APA regulation across a broad array of cell types and conditions, and the functional impact of 3’UTR identity, only a few proteins that regulate 3’ UTR isoform choice have been identified [1–3]. In this study, we create and describe a resource for the community to better understand the regulation of 3’UTR isoform expression by RBPs through processing available ENCODE data uniformly with DaPars. Our results provide a uniform platform to directly compare RBP impact on 3’UTR identity. Moreover, the results from our analysis can be directly compared with many other recent studies that utilize the DaPars software to categorize the breadth of 3’UTR isoform differences between tissues, brain disorders, immune cells, and cancers, and investigate how genetic variation can influence 3’UTR isoform expression in these cell types [10,11,21,44–48]. Such comparisons are expected to reveal putative regulators of physiologically-relevant APA regulation and motivate further research.

We acknowledge that the catalogue of 3’UTR regulators we describe here is somewhat limited by the fact that the transcriptomic data analyzed is only in HepG2 and K562 cells. However, we have previously demonstrated that analysis of RBP binding and functional targets of alternative splicing, another key pre-mRNA processing step, in model systems, including HepG2 and K562 data from ENCODE, can be used to learn regulatory relationships that are important in contexts beyond these model systems. For example, we previously described the antagonistic relationship between RBPs in the CELF and RBFOX families in heart and skeletal muscles or decreased QKI or PTBP1 expression in cerebellum controlling brain-specific splicing patterns based on similar cell line depletion and binding experiments, including from ENCODE [49,50]. This leads us to believe that our results for 3’UTR isoform expression can be applied to these broader contexts as well.

Other studies have set out to categorize regulators of alternative polyadenylation using single cell RNA-seq based approaches that leverage Perturb-seq techniques in order to query knockdown of tens to thousands of depletion experiments [51–53]. However, these genome wide perturbation experiments can be underpowered, requiring pooling of knockdown RBP experiments based on functional annotations [51]. Moreover, targeted perturbation experiments have examined a somewhat limited set of known factors [52]. Interestingly, DDX55 was identified in one of these studies on Perturb-seq in K562 cells as a part of a cluster of ribosome biogenesis genes that, when experiments are pooled together, regulate a number of 3’UTRs [51]. Therefore, our results complement these studies and expand the set of targets regulated by RBPs.

### DDX55 is a previously unrecognized direct regulator of 3’ UTR identity and may regulate APA at least in part via altering RNA secondary structure

Beyond providing a general catalog of RBPs that determine 3’ UTR identity and/or bind to 3’ UTRs, this study specifically identifies DDX55 as a direct regulator of 3’ UTR identity. The fact that DDX55 depletion leads to a similar set of 3’ UTR changes in both nuclear and cytoplasmic mRNAs (Figure 4), suggests that DDX55 regulates 3’UTR isoform expression primarily through choice of PAS in the nucleus. DDX55 is conserved down to yeast where it has previously been categorized as a regulator of rRNA biogenesis [35,36]. In human cells DDX55’s role in rRNA processing has recently been confirmed to bind domain IV of the 28S rRNA [54]. However, this study also showed DDX55 is mostly excluded from the nucleolus and sucrose gradients find DDX55 to be enriched in fractions containing free proteins or small complexes, suggesting it may play other roles in RNA processing [54]. One indication of additional activity of DDX55 was from a recent study that implicated direct interaction between DDX55 and BRD4 as upregulating β-catenin signaling in hepatocellular carcinoma [55]. While an effect on 3’UTR isoform expression was not tested in that study, BRD4 is known to interact with and recruit the cleavage and polyadenylation machinery during transcription elongation [56] and co-inhibition of BRD4 and TOP1 induces readthrough transcription with implications for other cancers [57,58]. In another work, DDX55 was found to co-IP with CTCF and cohesion, which have also been implicated in the regulation of alternative polyadenylation [59,60].

While the interaction of DDX55 with co-factors such as BRD4 and CTCF may explain some of DDX55’s activity in APA regulation, sequence analysis of DDX55-regulated PAS further suggests that DDX55 may carry out at least some of its APA regulatory activity by altering RNA structure in the 3’ UTR (Figure 5). Studies have shown that CFIm, which binds UGUA upstream of PAS, is dispensable for the cleavage and polyadenylation (CPA) reactions, but instead enhances assembly of the 3’-end processing complex and increases the reaction efficiency. Importantly, *in vitro* and *in vivo* assays have demonstrated CFIm acts in a position-dependent manner and enhances the CPA reaction most efficiently at −50 nt upstream of the cleavage site by enhancing recruitment of CPSF that recognizes the PAS hexamer. Our analysis of transcriptome-wide eCLIP signals of DDX55 suggests that DDX55 also preferentially binds around UGUA motifs in a region near transcript ends; however, the spacing of UGUA from the PAS hexamer, and thus the cleavage site, is significantly more distant compared to CFIm68. Moreover, we find that the long linker region between UGUA and the PAS hexamers near DDX55 binding sites have increased G/C content and predicted minimal free energy by *in silico* RNA folding compared with CFIm68.

Further supporting a role for DDX55 binding structured regions, recent deep learning models that predict RBP binding events that leverage CLIP, nucleotide sequence, and secondary structure data (PrismNet), found DDX55 tends to bind structured sequences [61,62]. Strikingly, DDX55 was among the worst-performing RBPs for predictions utilizing sequence alone and showed the most dramatic improvement when leveraging RNA structure information (AUC of sequence mode: 0.72, AUC of structure mode = 0.87). Furthermore, expression of full-length, recombinant DDX55 was found to bind double-stranded RNA with high affinity by fluorescence anisotropy [54].

In sum, our work, together with previous studies of DDX55, suggest several mechanisms by which DDX55 may directly influence PAS choice to regulate APA. We note, however, that we cannot fully exclude that DDXX55 may also have additional activities that control 3’UTR identity. Indeed, we have identified at least a few genes for which DDX55-dependent changes in 3’ UTR use were more apparent within cytoplasmic mRNA than nuclear mRNA, indicating a potential role in 3’ UTR-mediated mRNA stability.

## Conclusions

Alternative polyadenylation is a widespread gene regulatory mechanism that controls 3’ UTR identity and can influence mRNA and protein abundance and duration of expression. However, our understanding of protein factors that regulate 3’ UTR expression remains limited. Here we leverage data from hundreds of RBP depletion studies in the ENCODE database to identify factors that regulate 3’UTR identity, many of which have not previously been reported to impact gene expression in this way. We demonstrate the utility of our broad analysis by pursuing DDX55, and provide evidence that DDX55 is a bona fide regulator of APA by altering use of structured PAS elements. Our study thus identifies at least one previously unrecognized regulator of APA and opens the door to many new mechanistic studies for a host of additional RBPs.

## Methods

### RNA-seq data processing and definition of tandem 3’UTR PAS shifts

Raw fastq files were downloaded from the ENCODE portal (http://www.encodeproject.org). All RNA-seq samples used in this study are listed in Supplementary Table 1. Adapter sequences were removed and low quality ends were trimmed using BBDuk, part of the BBTools suite (http://sourceforge.net/projects/bbmap/), using the provided adapter fasta file and using the following parameters: `ktrim=r, k=23, mink=11, hdist=1, minlength=35, tpe, tbo, qtrim=r, trimq=20, qin=33`. Successful adapter removal and quality trimming was confirmed using FastQC.

Trimmed fastq files were aligned to hg38 genome using STAR (v2.7.1a) supplemented with splice junctions from Ensembl v94 and the option `--alignSJoverhangMin 8`. We retained uniquely mapped reads for downstream analysis.

To identify tandem PAS shifts within terminal exons we utilized DaPars (v0.9.1) [21] based on the hg38 RefSeq annotation from NCBI. We applied a coverage cut off to define expressed terminal exons using `Coverage_cutoff=30`. Changing terminal exons as those containing an absolute delta Percent Distal Usage Index (dPDUI) of 20% or more with an adjusted *p*-value of less than 0.05 when comparing RBP depletion samples with the appropriate controls (Supplementary Table 1). Non-changing terminal exons were defined as those with an absolute dPDUI < 5% and an adjusted *p*-value > 0.05. We found these thresholds previously to yield high-confidence predictions that experimentally validate in the context of CELF2 depletion [14]. We counted the number of unique genes that had at least one changing terminal exon and further categorized these into terminal exons where the RBP lengthens the 3’UTR (i.e. RBP promotes long 3’UTR expression or distal PAS usage resulting in a shift to the proximal PAS upon depletion) and terminal exons where the RBPs shortens the 3’UTR (promotes proximal PAS usage resulting in a shift to the distal PAS upon depletion). These results are summarized in Supplementary Table 2 for all RBPs in both cell lines.

### Updated eCLIP peak calling to include regions downstream of PAS

To quantify the effects of extending the annotated transcript/gene ends in order to capture CstF and, potentially, other RBP binding events in ENCODE, updated the provided gff annotation files CLIPper (v1.2.1) used in peak calling to include a downstream extension of 500 nucleotides. We downloaded pre-trimmed and aligned bam files from the ENCODE portal (Supplementary Table 3) and extracted read two using Samtools view requiring flag 128. We used CLIPper (v1.2.1) with the read two bam files as input with the following options to call peaks: `--bonferroni --superlocal --threshold-method binomial`. These updated peaks were used to identify PAS proximal binding proteins (see below).

### eCLIP peak analysis to identify 3’UTR and PAS proximal binding proteins

To identify RBPs that bind to expressed 3’UTRs we defined expressed transcripts based on the median expression across all control replicates from polyA selected libraries corresponding to shRNA depletion experiments. We quantified transcript expression for each control sample using Salmon (v0.14.1) [63] quasi-mapping on the trimmed fastq files based on Ensembl v94 transcriptome annotation indexed to k-mer length 31. Transcripts with a median expression ≥1 TPM across all controls in HepG2 or K562 cells were retained and we extracted 3’UTR coordinates and overlapped these regions with differing levels of eCLIP peak calls (experiments summarized in Supplementary Table 3). We defined an expressed 3’UTR to be bound by an RBP in a cell type if there was an overlapping eCLIP peak in either replicate eCLIP experiment within the same cell type. We then ranked the RBPs based on which had any binding evidence in the highest fraction of expressed 3’UTRs. To identify RBPs that bind specifically to 3’UTRs, we took high-confidence eCLIP peaks, based on ENCODE’s Irreproducible Discovery Rate (IDR), and overlapped them to expressed 3’UTR coordinates in the appropriate cell type. We ranked RBPs based on which IDR peak set showed the highest fraction of peaks that overlapped expressed 3’UTRs.

To identify RBPs that bind specifically near PAS and to extend our results to human tissues, we downloaded 3’end sequencing read counts from APASdb [64] corresponding to human liver and human lymph nodes. We defined highly expressed PAS as those with 100 or more reads in each tissue and overlapped those to downstream extended eCLIP peak calls (see above) for all RBPs within 500 nt of a highly expressed PAS. HepG2 eCLIP peak calls were overlapped with liver PAS and K562 peaks with lymph node PAS and plotted the per nucleotide frequency of overlap between PAS and binding for every RBP.

### Motif analysis

To discover the motifs DDX55 binds, we extracted the sequences underneath HepG2 DDX55 IDR peaks and used Homer [65] to find enriched motifs using `findMotifsGenome.pl` with the following parameters `-p 4 -rna -S 10 -len 4,5,6,7,8 -noknown`. When plotting per nucleotide motif frequencies around eCLIP peaks and DDX55 regulated PAS we searched for motif occurrence using a sliding window approach of 10 nucleotides for 4-mers associated with the CFIm motif (UGUA) and its reverse complements and 20 nucleotides for 6-mers associated with the PAS hexamers (AWUAAA) and its reverse complements. Frequencies were smoothed using a running mean of 5 nucleotides in these cases. For G/C and A/U content plots, frequencies at each position were smoothed using a running mean of 10 nucleotides.

### Cell culture and transfection

Low passage HepG2 cells were grown on 10 cm plates to 75% confluency in 10mL DMEM +10% FBS and transfected with 120 pmol of either DDX55 siRNA or a non-targeting pool of siRNA with 25 μL RNAiMAXusing the forward transfection protocol using Opti-MEM. This was repeated for three biological replicates (passages 2, 3, and 5). After 45-50 hours, cells were harvested and resuspended in 2 mL DPBS which was used for protein and RNA extraction. For protein fractionation cells were incubated on ice in 10mM HEPES pH 7.9, 1.5nM MgCl2 10mM KCL for 5 minutes and lysed on ice for 5 additional minutes with the addition of NP40 to 0.1%. Nuclei were collected by low speed centrifugation, washed three times with DPBS, and then lysed with RIPA buffer. For RNA fractionation cells were lysed on ice for 5 minutes in 10mM Tris-HCl pH 8.0, 0.32 sucrose, 3mM CaCl2, 2mM MgCl2, 0.1mM EDTA 0.1% NP40 1mM DTT plus 0.04U/ul RNasin. Nuclei were collected by low speed centrifugation, and Trizol was added to the supernatant for the cytoplasmic fraction or the pellet for the nuclear fraction.

### Western blotting

10 μg of whole cell protein was loaded onto an 8% gel and tank transferred for 1 hour at 100V at room temperature with chilled transfer buffer. Probed ∼14 hours at 4℃ with a 1:1,000 dilution of rabbit α-DDX55 antibody (Sigma HPA044101), or a 1:5,000 dilution of α-GAPDH antibody (Cell Signaling, 2118) or α-lamin B1 (Abcam, ab133741) for 1 hour at room temperature.

### 3’end sequencing and processing

Quality of total RNA from whole cell, nuclear, or cytoplasmic fractions from cells transfected with siRNA against DDX55 or a non-targeting pool control in triplicate was confirmed using a Bioanlyzer with RIN≥8.4 for all samples (range 8.4-9.4). Samples were submitted to Lexogen for QuantSeq REV V2 library preparation and sequencing to generate 76 nt, single end reads corresponding to the 3’end of polyadenylated RNAs. Cutadapt was used to remove adapter sequences and polyA or polyT sequences. Trimmed reads were aligned to the genome using STAR (v2.6.1a) with the following parameters --outFilterType BySJout --outFilterMultimapNmax 200 --outFilterMismatchNmax 999 --outFilterMismatchNoverLmax 0.6 --seedPerWindowNma× 10 --alignIntronMin 20 --alignIntronMa× 1000000 --alignMatesGapMa× 1000000 --alignSJoverhangMin 8 --alignSJDBoverhangMin 1. Samples were sequenced to a depth of 18.2 to 29.4 million reads with a uniquely mapped read percentage of ∼84-88%.

Because the 3’end of the read corresponds to the site of cleavage and polyadenylation and since the cleavage event is somewhat imprecise, we defined 3’end clusters to quantify site usage. To unify cluster definitions across all samples, we combined read coverage information from all 18 samples and created clusters by merging continuous regions with more than 10 reads at every position. We removed resulting clusters that had fewer than 20 reads within the clusters across all samples. The polyadenylation site (PAS) for each cluster was assigned to be the single nucleotide position with the highest read coverage within the cluster. In the event of multiple positions with the same coverage, the 3’ most position was chosen.

Internal priming from genomically encoded A-rich regions of transcripts is problematic for many targeted 3’end sequencing techniques. We addressed this using multiple heuristics to remove potential internal priming events. First, we removed clusters with more than 6 consecutive genomically encoded adenosines directly downstream of the defined PAS for the cluster. We also removed clusters that have more than 7 As within the 10 nt directly downstream of the cluster. Next we focused downstream analysis on clusters that contained one of the top two PAS hexamer sequences (AAUAAA or AUUAAA) within the cluster. Lastly, we required clusters to be present in PolyA_DB, a database of high-confidence PAS based on 3’READS techniques which avoids issues of internal priming [66].

After cluster definition and filtering, we focused on clusters that occurred within terminal exons and calculated relative usage of each cluster based on the total reads of that cluster in each sample over the sum of all clusters with the same terminal exon. We defined changes upon DDX55 knockdown by comparing each triplicate knockdown to its matched control experiment and found PAS clusters with changes of 10% or more in two out of three replicates in the same direction and no comparison in the opposite direction. Overlap of confident changing terminal exons in the nuclear and cytoplasmic compartments resulted in 201 shared terminal exons. To define compartment specific changes, we defined confident non-changing terminal exons in each as those that had a total of at least 10 reads in each sample with no PAS changing more than 5% in any replicate knockdown. This resulted in only 12 cytoplasmic specific and 3 nuclear specific terminal exons.

### 3’RACE

3’RACE was performed using the SMARTer RACE 5’/3’ Kit (Takara Biosciences) kit according to the manufacturer’s protocol and cDNA were analyzed on agarose gels. Gene specific forwarded primers used for amplification of 3’RACE cDNA are listed in Supplementary Table 4.

## Declarations

## Ethics approval and consent to participate

Not applicable

## Consent for publication

Not applicable

## Availability of data and materials

All supplementary data is publicly available. Processed data, including downstream extended eCLIP peak calls, and code to reproduce figures have will be published after peer review. Raw and processed 3’ end sequencing data have been deposited at GEO and will be available after peer review.

## Authors’ contributions

MRG, YB and KWL conceived of the project and wrote the manuscript. MRG carried out all of the data analysis under the direction of YB and KWL. TC and MJM carried out the experimental studies. All authors revised and approve of the manuscript.

## Acknowledgements

The authors thank members of the Lynch and Barash laboratories for useful comments and suggestions, and members of the APAeval team for helpful discussion.

