## Supplemental_Figures for "Integrative analysis of RNA binding proteins identifies DDX55 as a novel regulator of 3’UTR isoform diversity"

A

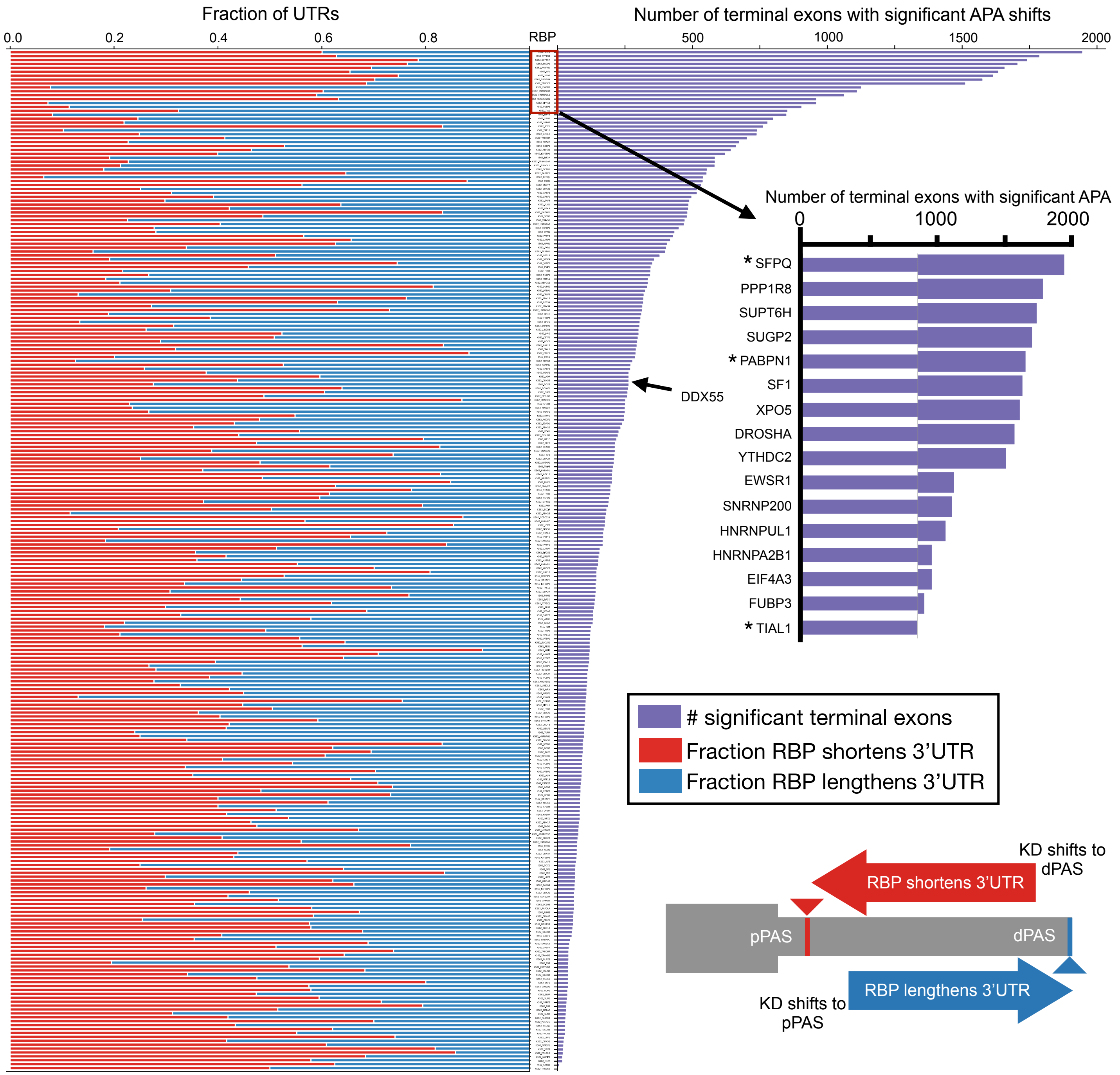


**Supplemental Figure 1**: **The landscape of 3'UTR tandem APA regulation in K562 cells**

**(A)** Summary of DaPars analysis of all K562 RBP knockdowns from ENCODE (244 shRNA mediated RBP depletion experiments) showing the number of significant terminal exons (right, purple, $|dDPUI| \geq 20\%$ with adjusted $p<0.05$). Right inset shows the top 16 RBPs by number of genes regulated. Stacked bar chart indicates the fraction of significant genes that had a pattern of RBP shortens 3'UTR (red, left bars) or RBP lengthens 3'UTR (blue, right bars) as described the inset cartoon. All experiments analyzed and counts are listed in Supplementary Tables 1 and 2.

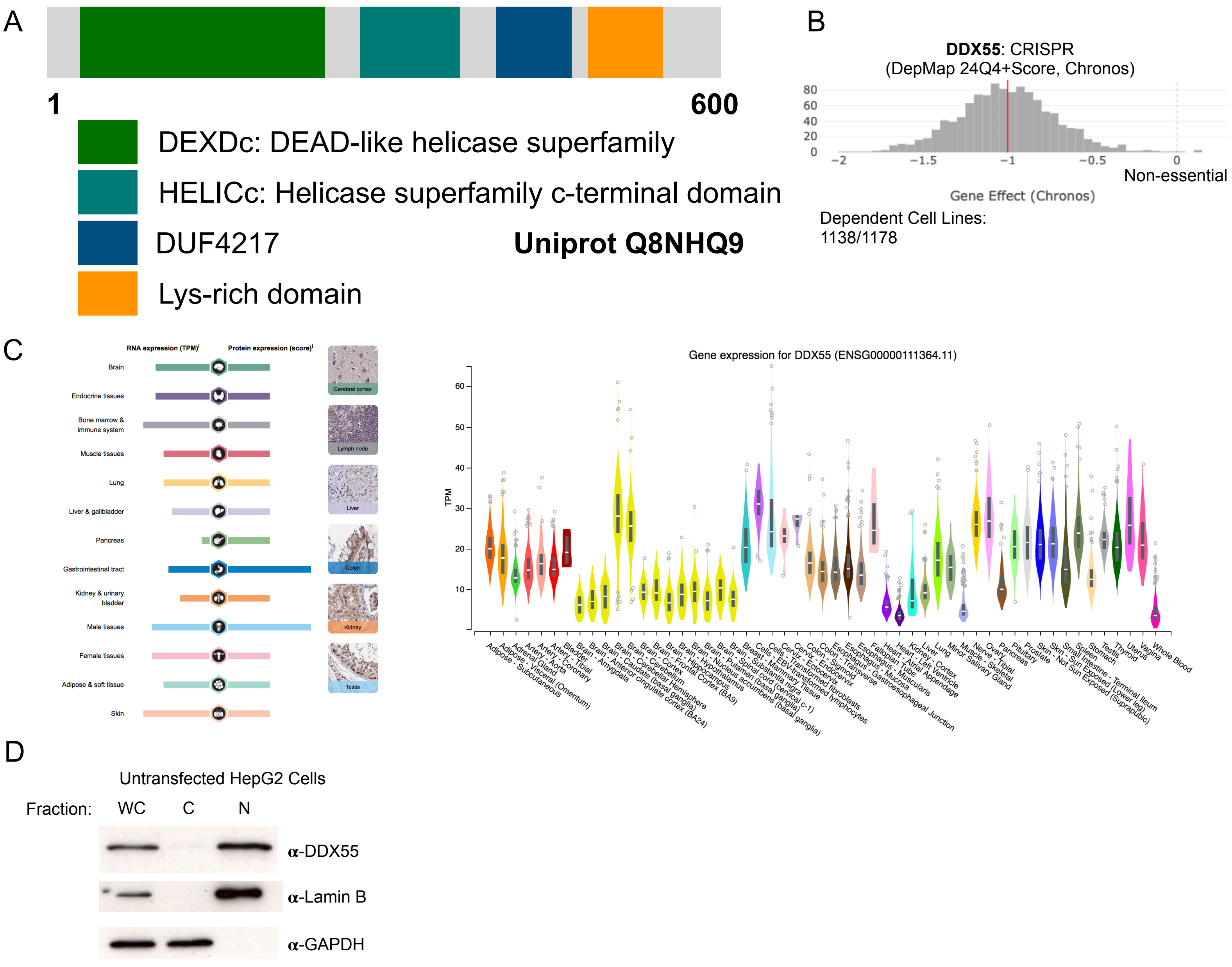


**Supplemental Figure 2**: **Characteristics and expression pattern of DDX55**

(**A**) Domain structure of DDX55 from Uniprot. (**B**) DepMap essential gene results showing gene effect prediction from CRISPR knockout in cell lines. Red line and score of -1 indicates the median value observed for pan-essential genes and 0 indicates non-essential. 1,138 out of 1,178 cell lines were found to be dependent on DDX55. (**C**) Human Protein Atlas summarizing RNA and protein expression of DDX55 across different tissues (left) and GTEx steady state mRNA expression level of DDX55 across indicated tissues. (**D**) Western blot on protein lysates from whole cell (WC), cytoplasmic (C), and nuclear (N) fractions from wildtype, untreated HepG2 cells. GAPDH and Lamin B served as controls for cytoplasmic and nuclear fractionation, respectively.

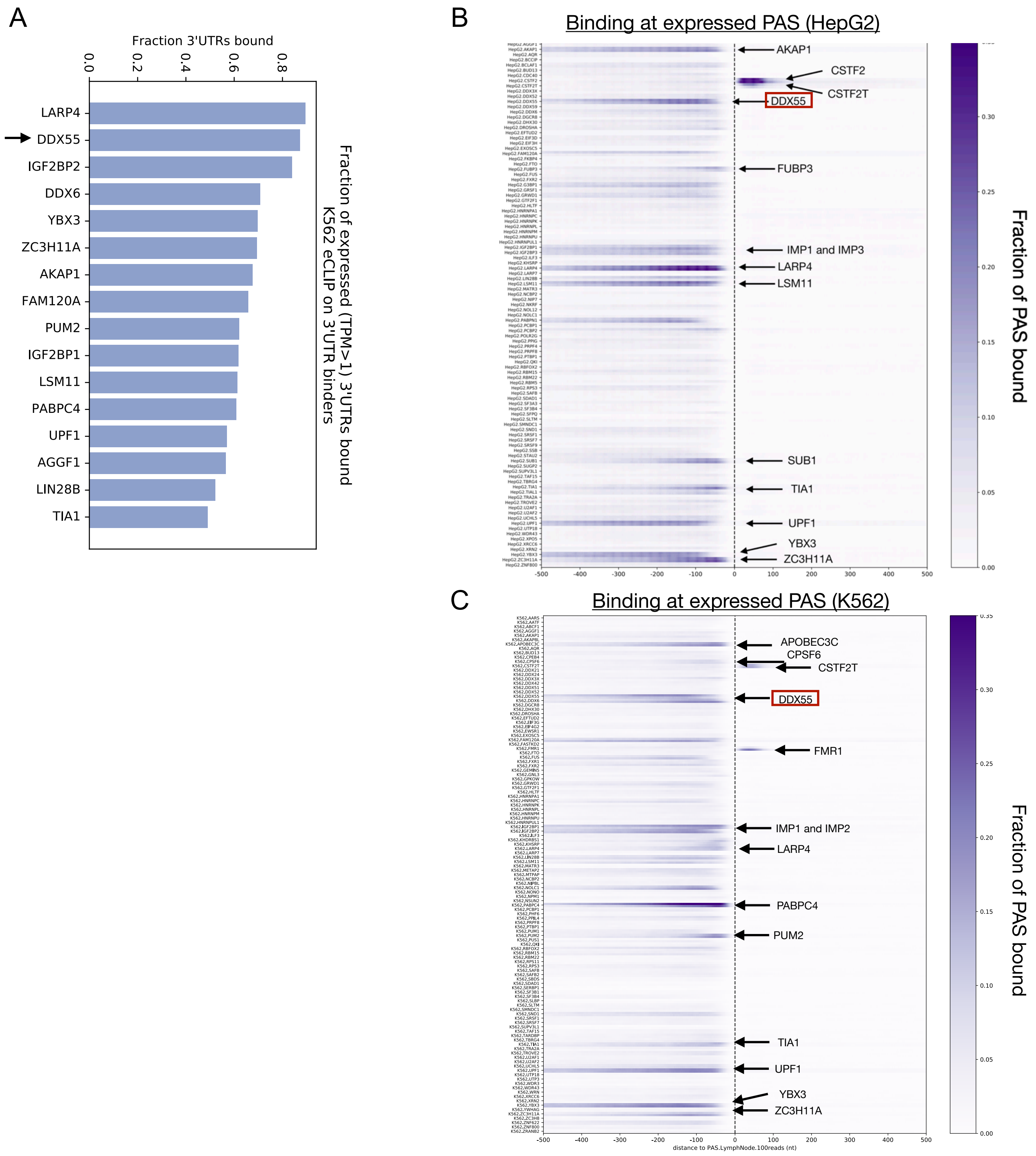


**Supplemental Figure 3**: **RBP eCLIP around 3' ends in K562 and HepG2 cells after extended peak calls**

(**A**) Fraction of expressed mRNA transcripts (TPM≥1) from K562 cells that had evidence of RBP binding in K562 cells (presence of any eCLIP peak) within the 3'UTR. (**B**) Heatmap showing the fraction of highly expressed PAS from human liver (100+ reads) that had evidence of proximal eCLIP binding (after downstream addition to peak calls) for all RBPs in HepG2 cells on a per-nucleotide basis within 500 nt up or downstream of the PAS. Select RBPs of interest are highlighted. (**C**) Same as panel (B) but for K562 cells. Select RBPs of interest are highlighted.

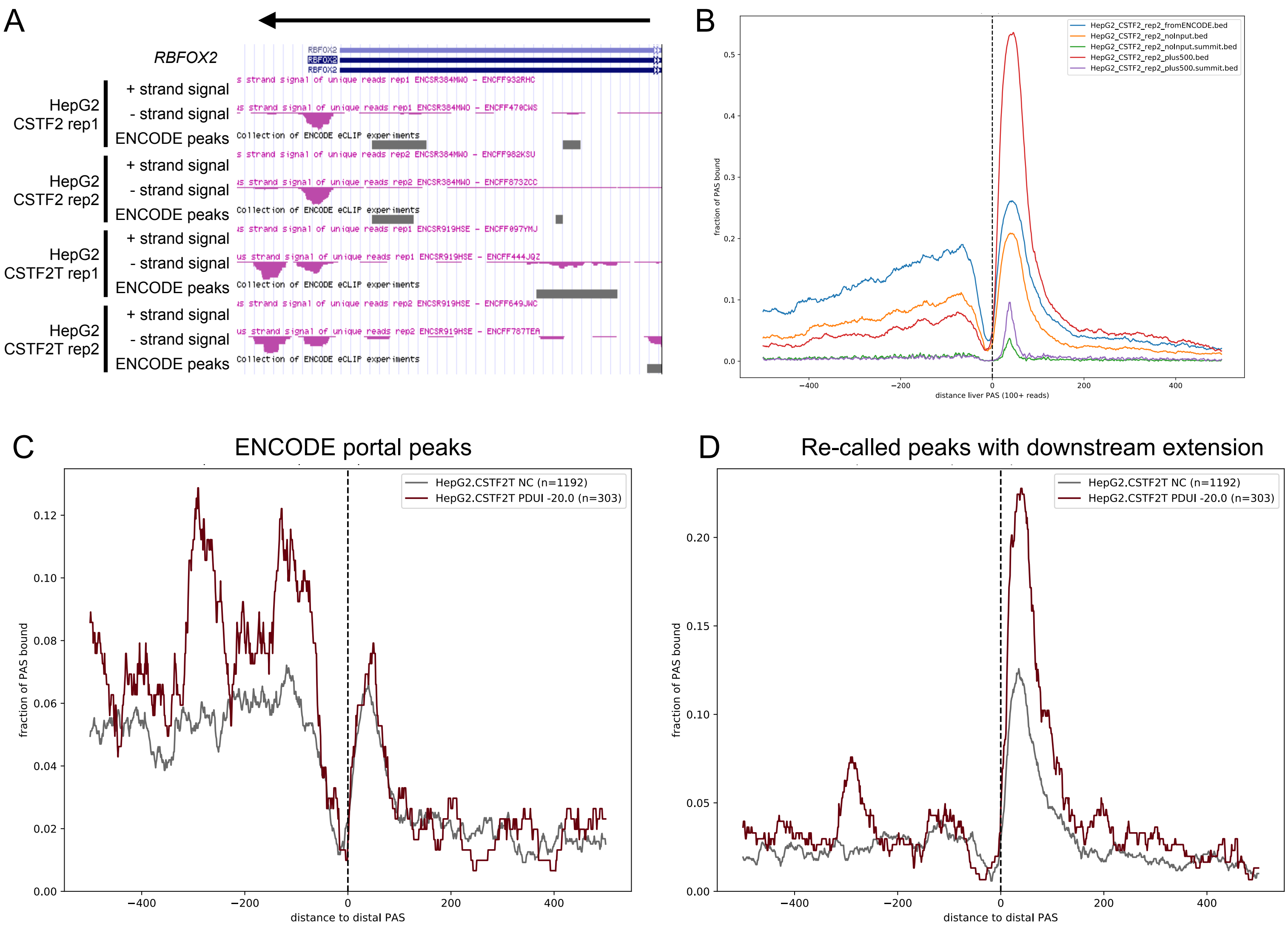


**Supplementary Figure 4: Correcting ENCODE eCLIP peak calling on RBPs that bind downstream of the terminal PAS**

(**A**) UCSC genome browser view of minus strand gene *RBFOX2*'s 3'end with ENCODE portal tracks for CSTF2 (CstF64) and CSTF2T (CstF64t) eCLIP read coverage from HepG2 cells (fuchsia). ENCODE called eCLIP peaks are shown in gray. (**B**) RNA map meta plot centered on highly expressed PAS from human liver (100+ reads) showing the fraction of PAS with HepG2 CSTF2 eCLIP peaks called by various methods labeled in the inset. (**C**) RNA map meta-transcript plot centered on the distal PAS showing the fraction of events with HepG2 CSTF2T eCLIP peaks called by ENCODE at each position around non-changing genes (gray) and genes where CSTF2T normally promotes transcript shortening (red). (**D**) Same as (C), but for HepG2 CSTF2T peaks re-called with a downstream extension of 500 nt to annotated transcript end.

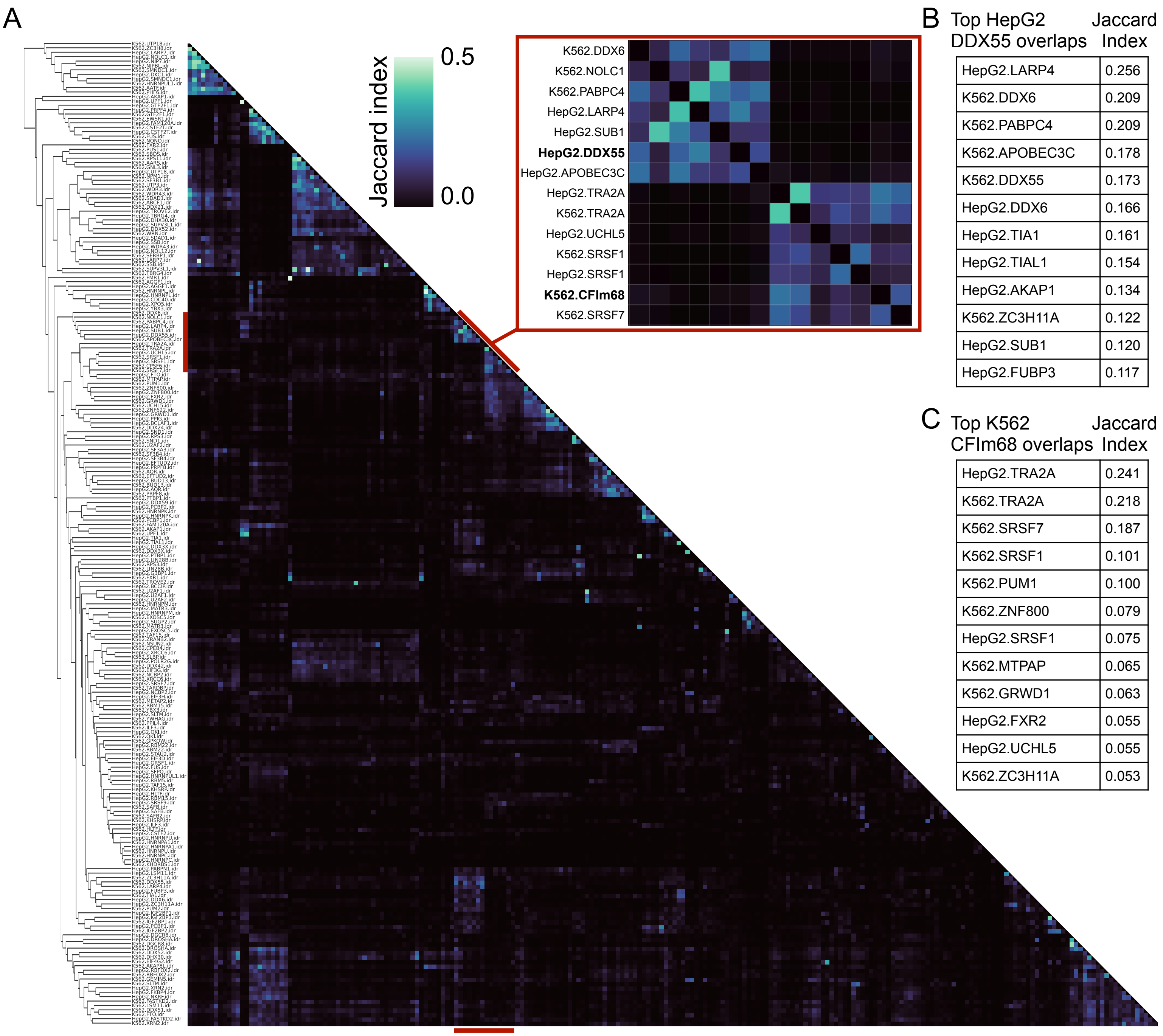


| Top HepG2 DDX55 overlaps | Jaccard Index |
|---|---|
| HepG2.LARP4 | 0.256 |
| K562.DDX6 | 0.209 |
| K562.PABPC4 | 0.209 |
| K562.APOBEC3C | 0.178 |
| K562.DDX55 | 0.173 |
| HepG2.DDX6 | 0.166 |
| HepG2.TIA1 | 0.161 |
| HepG2.TIAL1 | 0.154 |
| HepG2.AKAP1 | 0.134 |
| K562.ZC3H11A | 0.122 |
| HepG2.SUB1 | 0.120 |
| HepG2.FUBP3 | 0.117 |

| Top K562 CFIm68 overlaps | Jaccard Index |
|---|---|
| HepG2.TRA2A | 0.241 |
| K562.TRA2A | 0.218 |
| K562.SRSF7 | 0.187 |
| K562.SRSF1 | 0.101 |
| K562.PUM1 | 0.100 |
| K562.ZNF800 | 0.079 |
| HepG2.SRSF1 | 0.075 |
| K562.MTPAP | 0.065 |
| K562.GRWD1 | 0.063 |
| HepG2.FXR2 | 0.055 |
| HepG2.UCHL5 | 0.055 |
| K562.ZC3H11A | 0.053 |

**Supplementary Figure 5: Pairwise eCLIP overlaps**

(**A**) Heatmap showing hierarchical clustering of Jaccard Index for all pairwise IDR eCLIP peak sets. Inset highlights two clusters that contain DDX55 and the CFIm subunit CPSF6 (CFIm68). (**B**) Top 12 IDR peaks from eCLIP experiments indicated that overlap with HepG2 DDX55 IDR peaks, sorted by Jaccard index. (**C**) As in (B), but for overlaps with K562 CPSF6 (CFIm68) IDR peaks

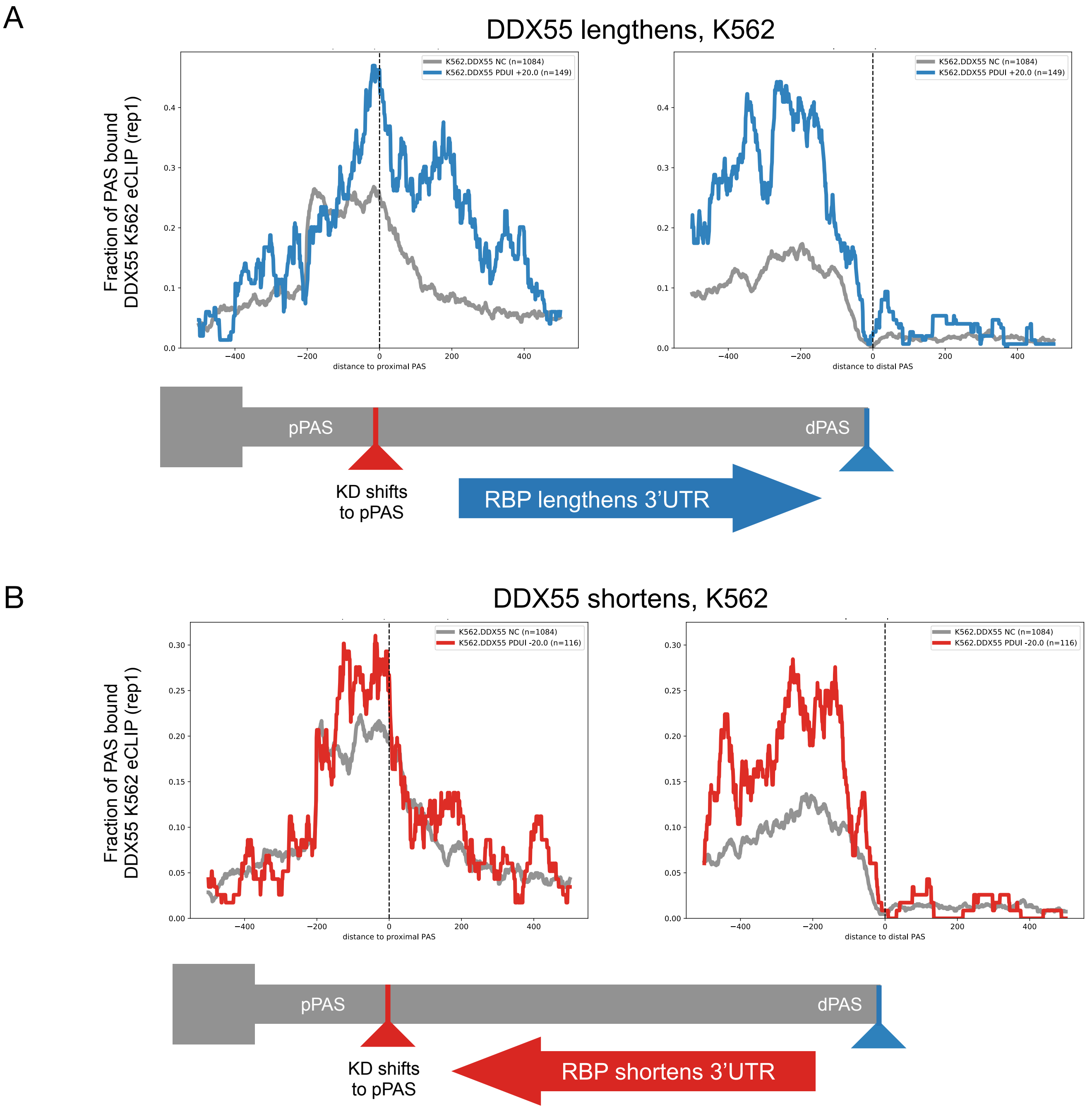


**Supplementary Figure 6**: **RNAmaps DDX55 tandem 3'UTR regulation in K562 cells**

(**A**) RNA map of the fraction of events with K562 DDX55 eCLIP peaks (replicate 1 shown) centered around proximal polyadenylation sites (pPAS, left) or distal PAS (dPAS, right) for events in K562 cells where DDX55 promotes 3'UTR long isoform expression (blue line, knockdown (KD) shifts expression towards pPAS, dDPUI≥20% with FDR<0.05) or events were DDX55 depletion had no effect on PAS choice (gray line, |dPDUI|≤5% with FDR>0.05). (**B**) same as panel (A) but for events in K562 cells where DDX55 promotes 3'UTR short isoform expression (red line, knockdown (KD) shifts expression towards dPAS, dDPUI≤-20% with FDR<0.05) or events were DDX55 depletion had no effect on PAS choice (gray line, |dPDUI|≤5% with FDR>0.05).

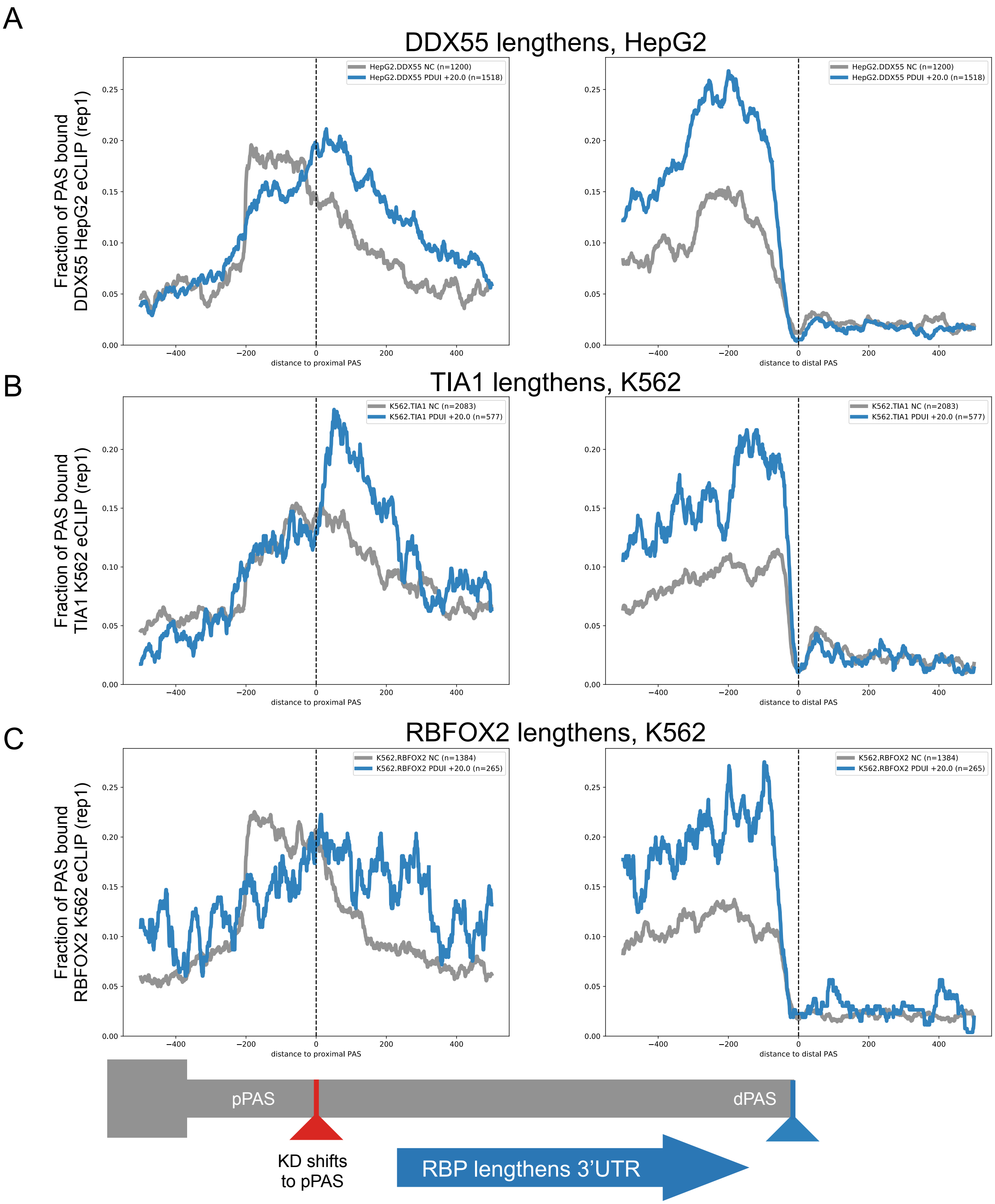


**Supplementary Figure 7**: **RNAmaps for putative direct regulators of promoting long 3’UTR isoform expression.**

(**A**) RNA map of the fraction of events with HepG2 DDX55 eCLIP peaks (replicate 1 shown) centered around proximal polyadenylation sites (pPAS, left) or distal PAS (dPAS, right) for events where DDX55 promotes 3’UTR long isoform expression (blue line, knockdown (KD) shifts expression towards pPAS, dDPUI≥20% with FDR<0.05) or events were DDX55 depletion had no effect on PAS choice (gray line, |dPDUI|≤5% with FDR>0.05). (**B**) As in (A), but for TIA1 knockdown RNA-seq and eCLIP in K562. (**C**) As in (A), but for RBFOX2 knockdown RNA-seq and eCLIP in K562.

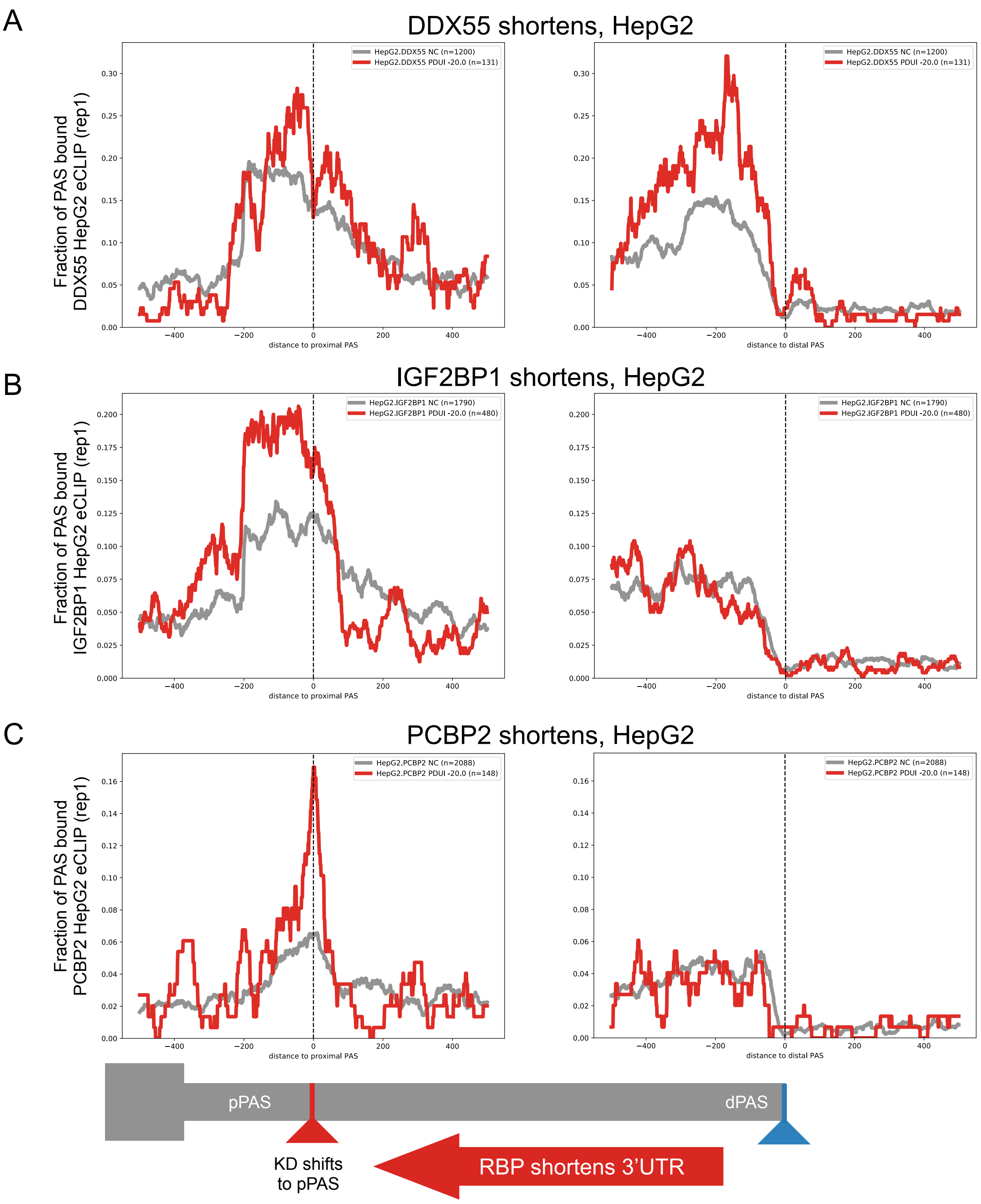


**Supplementary Figure 8: RNAmaps for putative direct regulators of promoting short 3'UTR isoform expression**

(**A**) RNA map of the fraction of events with HepG2 DDX55 eCLIP peaks (replicate 1 shown) centered around proximal polyadenylation sites (pPAS, left) or distal PAS (dPAS, right) for events where DDX55 promotes 3'UTR short isoform expression (red line, knockdown (KD) shifts expression towards dPAS, dDPUI≤-20% with FDR<0.05) or events were DDX55 depletion had no effect on PAS choice (gray line, |dPDUI|≤5% with FDR>0.05). (**B**) As in (A), but for IGF2BP1 knockdown RNA-seq and eCLIP in HepG2. (**C**) As in (A), but for PCBP2 knockdown RNA-seq and eCLIP in HepG2.

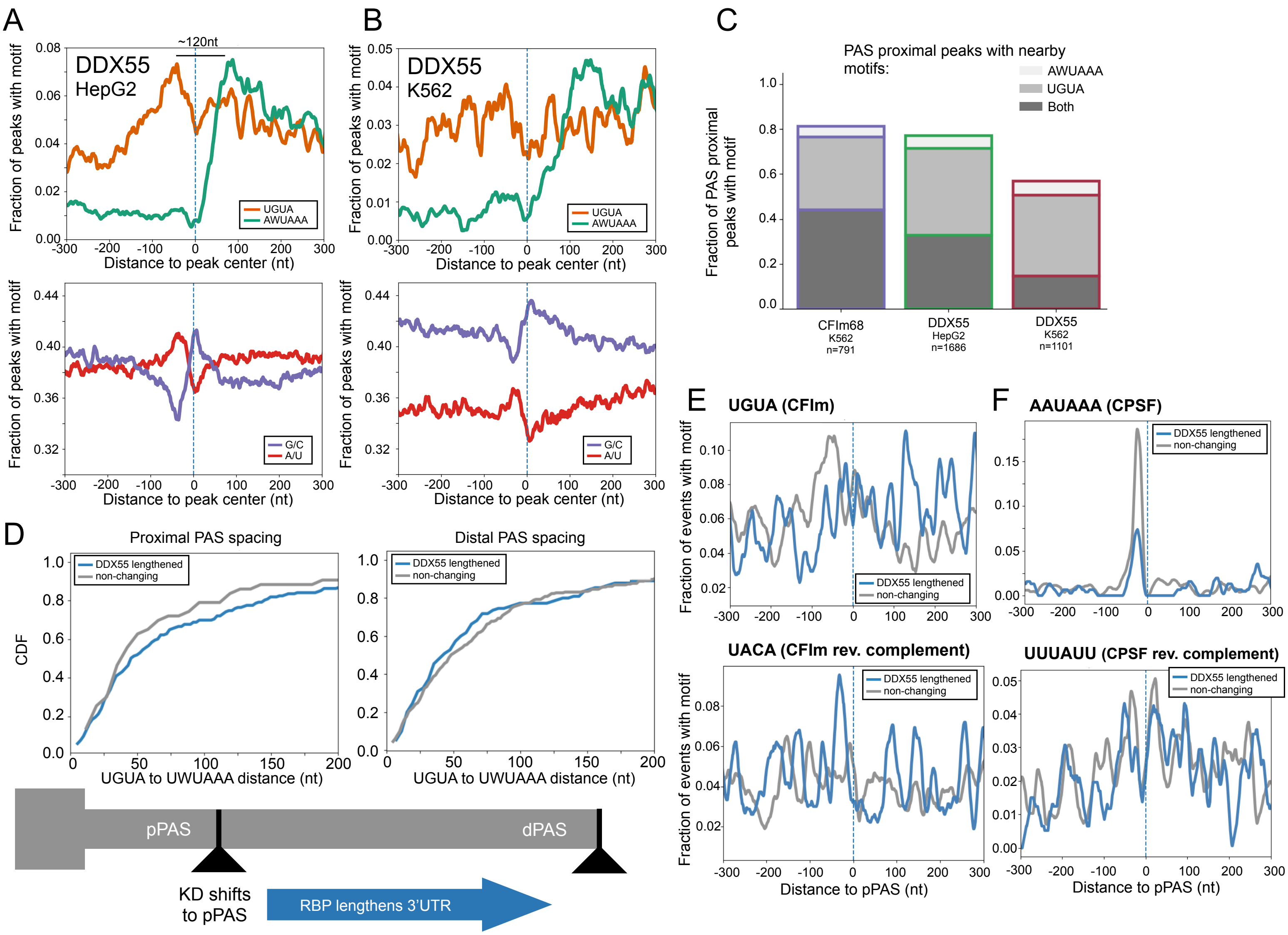


**Supplementary Figure 9: Additional sequence features of DDX55 and CFIm binding.**

(**A**) Motif maps showing per nucleotide frequency of the core PAS hexamers (AWUAAA, green) and CFIm upstream enhancing element (UGUA, orange) centered on the high confidence eCLIP peaks for DDX55 in HepG2 cells (top). Bottom motif map shows frequency of A/U (red) or G/C (purple) nucleotides over the same region centered on DDX55 peaks. Smoothing over a 10 nt window was applied. (**B**) As in (A), but for DDX55 high confidence eCLIP peaks in K562 cells. (**C**) Stacked bar chart showing fraction of subset of high confidence peaks for CFIm68 (CPSF6) or DDX55 near experimentally determined PAS which had one of the top two PAS hexamers alone (AWUAAA, light gray), the CFIm motif alone (UGUA, medium gray), or both (dark gray) within 150 nt of peak centers. (**D**) CDF of UGUA to AWUAAA motif distance occurring proximal to pPAS (left) and dPAS (right) that are regulated by DDX55 (blue, promotes 3'UTR lengthening) or non-regulated (gray). (**E**) Motif maps showing per nucleotide frequencies of the CFIm binding site (UGUA, top) or its reverse complement (UACA, bottom) around proximal PAS (pPAS) regulated by DDX55 (blue) or non-changing (gray). (**F**) As in (E), but for CPSF PAS hexamer (AAUAAA, top) or its reverse complement (UUUAUU, bottom) around proximal PAS.
